# Long-distance communication can enable collective migration in a dynamic seascape

**DOI:** 10.1101/2023.11.17.567622

**Authors:** Stephanie Dodson, William K. Oestreich, Matthew S. Savoca, Elliott L. Hazen, Steven J. Bograd, John P. Ryan, Jerome Fiechter, Briana Abrahms

## Abstract

Social information is predicted to enhance migratory performance, but the relative benefits of social information in the long-range movements of marine megafauna are unknown. In particular, whether and how migrants use nonlocal information gained through social communication at the extreme spatial scale of oceanic ecosystems remains unclear. Here we combine long-term acoustic recordings of foraging and migratory blue whales, oceanographic and prey data, and individual-based modeling to discern the cues underlying timing of blue whales’ breeding migration in the Northeast Pacific. We find that individual whales rely on both personal and social sources of information about forage availability in deciding when to depart from their vast and dynamic foraging habitat and initiate breeding migration. Empirical patterns of migratory phenology can only be reproduced by models in which individuals use long-distance social information about conspecifics’ behavioral state, which is known to be encoded in the patterning of their widely-propagating songs. Further, social communication improves pre-migration seasonal foraging performance by over 60% relative to asocial movement mechanisms. Our results suggest that long-range communication enhances the perceptual ranges of migrating whales beyond that of any individual, resulting in increased foraging performance. These findings indicate the value of nonlocal social information in an oceanic migrant and highlight the importance of long-distance acoustic communication in the collective migration of wide-ranging marine megafauna.

## INTRODUCTION

Across the animal kingdom, diverse taxa undertake long-distance movements to track the availability of resources in space and time (Abrahms *et al*. 2021a). The ability of migrating animals to match their timing and locations with optimal resource availability has critical consequences for individual energy gain, fitness, and population persistence (Both *et al*. 2006, Merkle *et al*. 2022, Middleton *et al*. 2018). The sources of information that animals rely on to make these movement decisions can determine how migratory populations respond to environmental variability (Winkler *et al*. 2014), how they find and exploit prey resources (Visser & Gienapp 2019), and how they fare under rapid environmental change (Post & Forchhammer 2007, Shaw 2016). As a result, considerable research has sought to understand the cues that animals use to decide when and where to move and migrate (Winkler *et al*. 2014; Berdahl *et al*. 2018; Aikens *et al*. 2022; Oestreich *et al*. 2022a). Nevertheless, which mechanisms guide long-distance migratory behavior remains a key question in ecology and this understanding is critical for anticipating the impacts of rapid environmental change on threatened migratory populations.

A range of cues, both internal and external and spanning fixed to dynamic, influence migratory decisions (Winkler *et al*. 2014, Shaw & Couzin, 2013). Many migratory animals rely on a range of information sources acquired via personal experience (Dall *et al*. 2005, Fagan *et al*. 2017, Abrahms *et al*. 2019) and/or social cues (Mueller *et al*. 2013, Jesmer *et al*. 2018 Aikens *et al*. 2022, Oestreich *et al*. 2022a). The relative utility of these information sources varies with life history traits, habitat complexity and predictability, and social context. For example, long-lived species may rely heavily on memory and learning gained through individual experience (Abrahms *et al*. 2019, Abrahms *et al*. 2021b, Polansky *et al*. 2015). Moreover, the relative utility of personal information depends on perceptual range, as nonlocal information allows the assessment of environmental variation over broader spatial scale in heterogeneous environments (Fagan *et al*. 2017).

Beyond individual sensing and learning, social information can play an important role in migratory decisions (Jesmer et al. 2018; Aikens *et al*. 2022; Oestreich *et al*. 2022a). This social information – including both intentional signals and inadvertent cues among individuals (Danchin *et al*. 2004; Dall *et al*. 2005) – can greatly enhance access to nonlocal information by expanding perceptual ranges well beyond that of any individual alone (Berdahl *et al*. 2013). Such information transfer can enable individuals to detect noisy environmental gradients and make better-informed migratory decisions, improving the ability to collectively track favorable resources and avoid predation risk in migration (Berdahl *et al*. 2018; Couzin 2018; Oestreich *et al*. 2022a). While such collective migrations are increasingly recognized throughout the animal kingdom, theory on emergent sensing and collective migration has primarily been developed and tested via study of social species which migrate in groups (e.g., salmon, Berdahl *et al*. 2017; and storks, Flack *et al*. 2018). Such theory has highlighted the importance of group size (King & Cowlishaw 2007; Kao & Couzin 2014) and density (Berdahl *et al*. 2017; Makris *et al*. 2009) in allowing for emergent sensing and collective behaviors including migration. Yet many species can communicate over substantial distance, indicating that collective migrations might emerge even in low-density populations (Couzin *et al*. 2018) and enable collective sensing of nonlocal information over even broader spatial scales than possible by aggregated groups. Theoretical simulations suggest that access to such nonlocal information should provide particular benefit in systems characterized by steep resource gradients and ephemeral resource patches (Fagan *et al*. 2017), yet tests of this hypothesis in the context of migration remain sparse.

The pelagic ocean represents roughly 99% of Earth’s habitable space by volume, and is characterized by dynamism and patchiness across spatial and temporal scales (Marquet *et al*. 1993, Steele 1991). In dynamic oceanic ecosystems, resources are extremely non-uniformly distributed, aggregating in ephemeral hotspots (Bertrand *et al*. 2014, Hazen *et al*. 2013). These attributes suggest that nonlocal information should provide particularly valuable information for pelagic animals’ long-distance foraging and migratory movements, yet whether and how migrants use nonlocal social information at the extreme spatial scale of oceanic ecosystems is unclear. Making detailed and persistent behavioral observations across ecological scales in vast and largely opaque oceanic ecosystems has been challenging historically, and limited the ability to test whether nonlocal social information gives rise to collective migrations in the pelagic. We address this question using multi-year, continuous empirical observations of population-level foraging and migration timing in blue whales (*Balaenoptera musculus*) in comparison to empirically-parameterized models of prey distribution, oceanographic conditions, and individual whale movement and communication.

Blue whales are obligatory krill predators and require significant and reliable prey resources to support their enormous body size (Goldbogen *et al*. 2019). In the Northeast Pacific Ocean, blue whales migrate in low densities between seasonally-productive foraging grounds off the west coast of North America in summer and fall, and calving grounds in the Gulf of California and west of Central America in the Costa Rica Dome during winter and spring (Bailey *et al*. 2009). The summer-fall foraging grounds represent the primary source of annual energy acquisition for individuals in this population (Pirotta *et al*. 2018). While these foraging grounds are in the consistently productive California Current system, they display considerable intra- and inter-annual variation. Within the foraging season, blue whales track their obligate krill prey that aggregate in ephemeral oceanographic habitat such as regions of enhanced upwelling or surface convergence (Fahlbusch *et al*. 2022; Ryan *et al*. 2022; Cade *et al*. 2021; Benoit-Bird *et al*. 2019). These fine-scale resource tracking strategies allow an adult blue whale to consume ∼1500 tonnes of krill during the foraging season in the California Current (Savoca *et al*. 2021), which fuels their long-distance migration to and from calving grounds and rearing of young (Pirotta *et al*. 2018).

At broader spatiotemporal scales, it is unknown how blue whales decide when to depart from these foraging grounds. Previous research has documented that blue whales track the long-term average ecosystem phenology as they migrate northward towards the feeding grounds (Abrahms *et al*. 2019). This strategy of reliance on memory over contemporaneous environmental cues maximizes their likelihood of energy acquisition in the long term (between years and over decades). At finer spatiotemporal scales, high-quality prey patches can be highly-dynamic (Abrahms *et al*. 2019). On their southward migration to breeding grounds, blue whales instead track contemporaneous ecosystem phenology, shifting the population-level timing of migration by as much as four months to match year-to-year variation in their foraging habitat (Oestreich *et al*. 2022b). But what signals and cues enable this plasticity in foraging and migration timing in such a vast and dynamic habitat?

Detecting the change in behavioral state of this blue whale population, from foraging to migration, is possible by evaluating the diel patterns of song production at an ecosystem scale (Oestreich *et al*. 2020). During the foraging months, blue whales in this population produce song primarily at night due to intensive daytime foraging effort, but transition to a more even diel song distribution following the cessation of foraging and onset of breeding migration (Oestreich *et al*. 2020). These diel patterns enable researchers to track population-wide behavioral transitions throughout the Northeast Pacific (Oestreich *et al*. 2022b; Pearson *et al*. 2023), thus prompting the question of whether blue whales themselves use long-distance information about conspecifics’ behavioral state to inform their migratory decisions.

Individual-based models (IBMs) are a modeling framework that can be used to examine how individual-based movement mechanisms generate population-level patterns. In this study, we encode a series of IBMs to investigate how population-level breeding migration behaviors emerge from hypothesized short-term foraging and migration decisions based on recent personal foraging experiences and/or social information gained from long-distance communication (Figure 1). We compare empirically-observed patterns of population-level migratory behavior to those emerging from the IBMs to deduce which decision mechanisms best replicate empirical observations. Using this integrative framework, we assess two related hypotheses (Figure 1D) concerning *(i)* the information sources on which whales rely in deciding when to depart from foraging habitat and begin their southward migration; and *(ii)* the consequences of reliance on different cue types for foraging performance. The results of this approach yield insight on the cues underlying blue whales’ migratory decisions. More broadly, this approach elucidates the dynamics of collective migration in this widely-distributed, low-density population, a common space-use pattern for populations across the animal kingdom.

**Figure 1:**
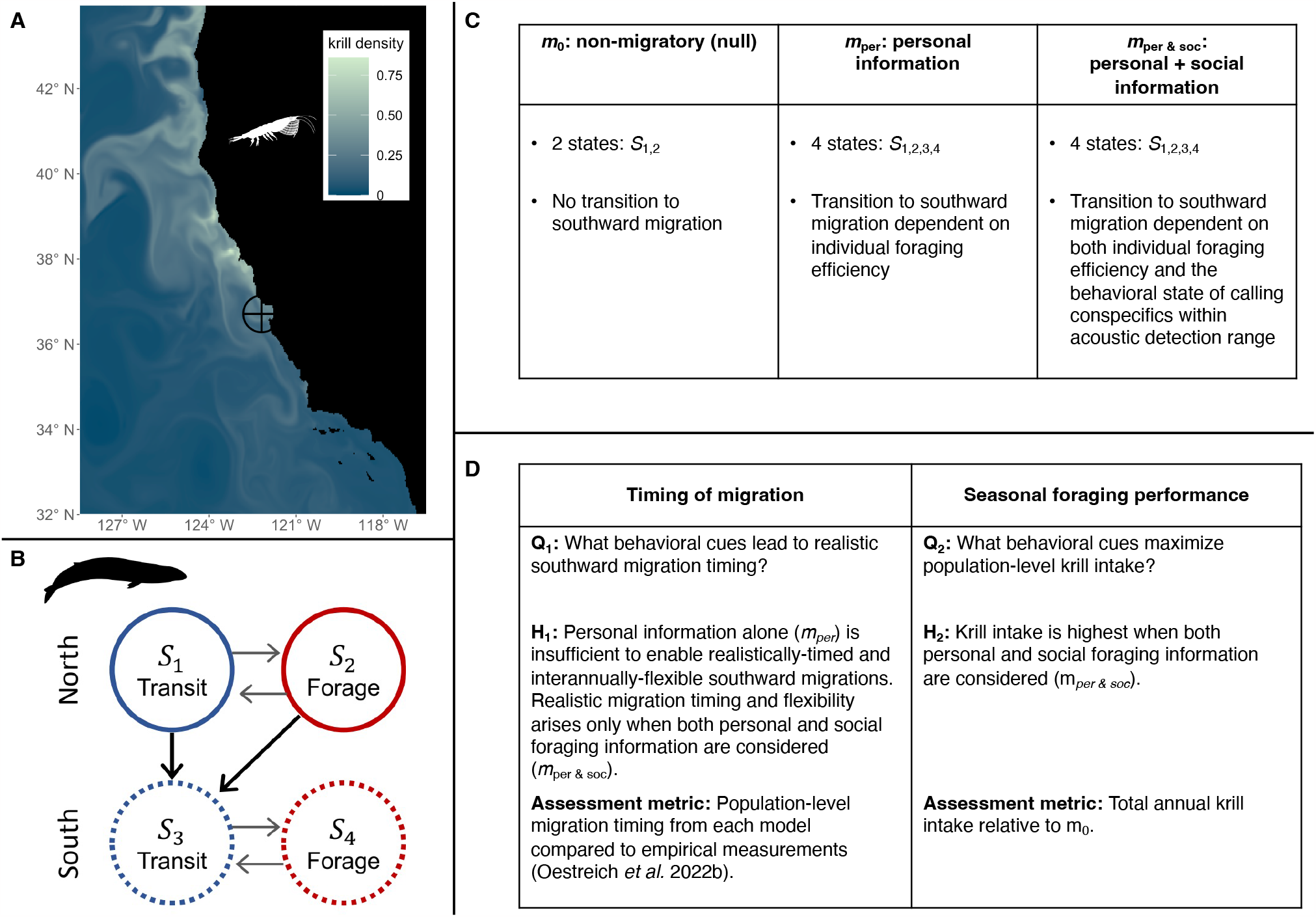
Model schematic. (A) Example of simulated near-surface krill density over the ROMS domain on yearday 250 (2004). The black circle indicates the approximate blue whale song detection range of the Monterey Bay hydrophone. (B) Behavioral states in the IBM. Arrows indicate possible behavioral state transitions. (C) Summary of tested model behavioral states and factors included in southward migration state transition function. (D) Summary of metrics used to assess questions and hypothesis.

## METHODS

### Environmental and Prey Data

Environmental conditions, specifically sea surface temperature (SST) and near-surface krill abundance, are provided from a Regional Ocean Modeling System (ROMS) implementation that has been coupled with a biogeochemical model, NEMUCSC, adapted from the North Pacific Ecosystem Model for Understanding Regional Oceanography of Kishi et al. (2007) and specifically parameterized for the California Current region (Fiechter *et al*. 2018; Fiechter *et al*. 2020). The model provides 3 km spatial and daily temporal resolution of SST and krill concentrations over a study domain of 116 – 128 °W and 32 – 44 °N for the years 1990 – 2010 (Fiechter *et al*. 2020); simulated krill concentrations have been evaluated against existing in situ data for May-June and are expected to adequately reproduce observed krill aggregation regions during the upwelling season (Fiechter *et al*. 2020). The ROMS-NEMUCSC output is pre-computed and supplied as input to the IBMs.

### Hydrophone Data and empirical social communications

Empirical studies (Oestreich *et al*. 2020; Oestreich *et al*. 2022b) found that blue whales’ population-level song call production increases throughout the foraging season in the Northeast Pacific, typically peaking from October through November. Only male blue whales have been recorded producing song calls in the Northeast Pacific (McDonald *et al*. 2001). Furthermore, the proportion of song produced during the night versus the day generally increases through at least September, then shows a drop toward a more even diel distribution of song. The timing of this drop in the proportion of song produced at night varies interannually (September – December; Oestreich *et al*. 2022b) and has been attributed to the population-level transition from foraging to southward migration (Oestreich *et al*. 2020). Because Northeast Pacific blue whales forage primarily during the day and foraging and singing behavior is temporally separate at sub-daily scale, predominantly nighttime singing behavior is strongly correlated with foraging behavior. As a result, the 24-hour patterning of blue whale song effectively provides long-distance information on the behavioral state of conspecifics in a vast and dynamic foraging arena (Oestreich *et al*. 2020). While the precise range over which blue whales can detect conspecifics’ songs remains uncertain, hydrophones can detect these songs at high signal-to-noise ratio over ranges exceeding 100 km (Oestreich *et al*. 2020). CT scanning and modeling-based efforts quantifying closely-related fin whale (*B. physalus*) hearing sensitivity (Cranford & Krysi, 2015) indicate auditory sensitivity of ∼100 dB re 1μPa in the frequency range of blue whale songs, which would allow for blue whale song detection over at least tens of km in Monterey Bay, CA, USA (Oestreich *et al*. 2020).

Data from the Monterey Accelerated Research System hydrophone (hereafter “Monterey Bay hydrophone”; Oestreich *et al*. 2020, Oestreich *et al*. 2022b) and tag analysis (Irvine *et al*. 2014) reveal median blue whale southward breeding migration dates generally fall in October through early November. High year-to-year variation is present with median departure dates ranging from mid-September to mid-December. Migrations for individual blue whales are recorded as early as late-July to as late as January (Oestreich *et al*. 2022b; Irvine *et al*. 2014).

### Individual-Based Model Framework

Individual-based models (IBMs) are a flexible model type that treat individuals as autonomous agents who follow a set of individual-level decisions and updates. Thus, IBMs are well suited to test how various strategies at the individual-level develop into population-level emergent behaviors, such as migrations. Comparing population-level behaviors of IBMs with empirical data provides an insight into the short-term decisions that lead to the broad-scale behaviors. IBMs have previously been used to understand the spatiotemporal movement of populations, including migrations of pelagic fish (Barbaro *et al*. 2009), resource-driven formation of locust bands (Bernoff *et al*. 2020), and group formation of zebrafish during movement (Oscar *et al*. 2023).

Our IBM is formatted as a probabilistic *N*-state switching model that treats individual whales as autonomous agents. Each state *S*_*k*_ captures known behavioral patterns and is associated with a distinct movement update. Previous analysis of tagging data revealed and parameterized distinct short-term movement distributions in blue whales associated with transit and forage behaviors (Bailey *et al*. 2009) throughout the foraging season. Furthermore, blue whales have been observed to definitively switch into southward migration behaviors, marked by an end in foraging and onset of intentional southward movement (Oestreich *et al*. 2020), with occasional pauses to forage. We use four behavioral states to characterize the distinct behaviors of the foraging and breeding migrations (Figure 1B). States *S*_*1,2*_ represent transit and foraging behaviors, respectively, during the northward foraging migration and *S*_*3,4*_ are transit and forage states for the southward breeding migration.

Dodson *et al*. (2020) presented a series of IBMs with state transitions based on sea-surface temperature (SST) and krill densities. These models accurately captured early-season foraging patterns of North Pacific blue whales but lacked inter-agent communication and models that included southward migration exhibited unrealistically early timings. The IBM presented here is an adaptation and extension of those parameterized in Dodson *et al*. (2020) with the goal of focusing on southward migration strategies. We utilize the previously defined transit-forage decisions (based on SST and krill) that allow within migration transitions (transitions from *S*_*1*_ to *S*_*2*_ and *S*_*3*_ to *S*_*4*_) and replace the inter-migration transitions that initiate the southward breeding migration (transitions from *S*_*1,2*_ to *S*_*3*_).

We provide a brief overview of the model here; full model details are available in the Supplement, including a description in the overview, design concepts, and details (ODD) framework, a protocol for standardizing published descriptions of IBMs (Grimm *et al*. 2010; Grimm *et al*. 2006). To focus on factors driving the breeding migration, yearly models are initiated on July 1^st^ with agents’ initial locations selected uniformly at random from areas of climatologically high krill densities from the ROMS-NEMUCSC solution (Abrahms *et al*. 2019). At every time-step (six-hour intervals), each agent uses a combination of local forage conditions (from ROMS-NEMUCSC), personal foraging experiences, and received social information to select a behavioral state with probabilities from a state transition matrix. A movement update (turning angle and step length) is selected from the designated behavioral state movement distribution, and the agent’s position is updated. Throughout the description, a subscript *n* indicates variables and quantities specific for agent *n*. Time dependence is indicated with *t*, corresponding to the time step.

### Model Implementation of Social Communication

Inter-agent communication is implemented by dividing communication into the subprocesses of call production and reception. Half of the agents are randomly initialized as male song producers. A proportion of male agents are randomly selected to call at each time-step. The proportion of calling male agents increases linearly throughout the season from 5 to 30%, in alignment with the increase in call intensities detected by the Monterey Bay hydrophone (Oestreich *et al*. 2020). The IBM uses daily mean simulated prey and SST data and does not distinguish night versus day. Instead, we define 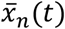 to be the average proportion of time spent foraging of agent *n* during the past 24 hours and define the call signal by 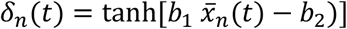 The call signal δ_*n*_*(t)* transforms 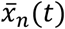 into a value between -1 and 1, with positive (negative) values indicating individual *n* spent more time foraging (transiting) during the time-period. The sign and value of δ_*n*_*(t)* is intended to mimic the diel patterns in hydrophone recordings of call intensities.

All agents act as call receivers. Received social information σ_*n*_*(t)* averages call signals from nearby individuals weighted with a distance-dependent amplitude that decays following the inverse-square law. In default parameter settings, call ranges are capped at a maximum distance of 125 km but maximum radii of 0 to 500km are explored (Figure 4). The sign of σ_*n*_*(t)* reflects the foraging behaviors of neighboring calling individuals and magnitude integrates the distance to calling whales with the frequency of foraging behaviors.

### State Transition Matrix and Southward Migration Decisions

Figure 1B defines the possible transitions between the four behavioral states and transition probabilities are dictated by elements of the state transition matrix τ

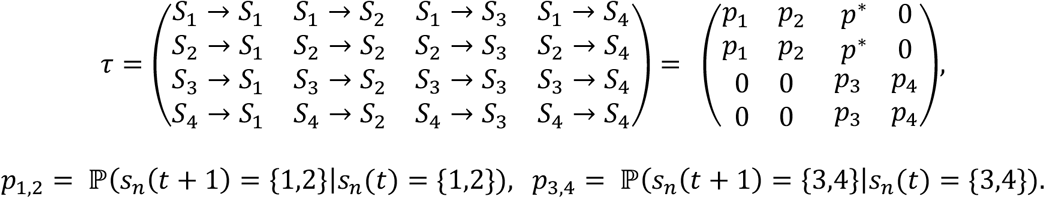

Zero elements of the state transition matrix represent unallowed transitions and *s*_*n*_*(t)* indicates the agent’s behavioral state. Southward migration decisions are incorporated into the model via the state transition matrix, specifically in the elements denoted by *p*^*\**^ that govern the transition probabilities from *S*_*1,2*_ to *S*_*3*_. The remaining transition probabilities are dependent on SST and krill values; these functions are given in the Supplement.

We test a suite of migration strategies, based on a combination of agents personal average foraging effectiveness 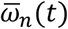 and average received social information 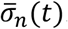 Foraging effectiveness 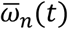 is defined as the average krill intake over a period of *T*-timesteps and gives a time-dependent measure of an individual’s personal foraging success. Foraging effectiveness and social information are averaged to allow individuals to explore their local environment and receive calls from multiple agents.

The following migration strategies prescribe different transition functions for *p*^*\**^ based on 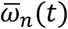and 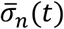

1. Individual foraging efficiency (personal, *m*_per_ (Figure 1C))

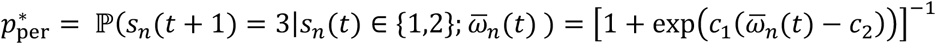
2. Individual foraging efficiency and social communication (personal & social, *m*_per&soc_)

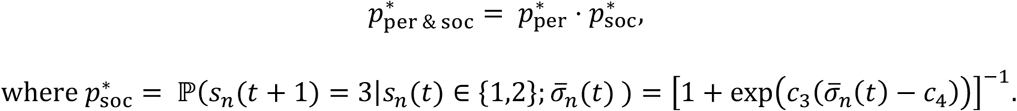

Values of constants are given in Supplementary Table 2. The personal strategy represents a mechanism based solely on an agent’s foraging efficiency. The multiplicative transition probability for the personal & social strategy reflects a strategy where agents initiate migration when both personal and social factors indicate poor foraging opportunities. A minimum krill intake requirement was explored as an additional asocial migration mechanism; a summary and discussion are included in the Supplement.

### Null Model

The results of the migratory models are tested against a hypothetical non-migratory null population (*m*_1_, Figure 1C). These agents follow the two-state transit-forage model of Dodson et al. (2020) and do not receive any southward migration cues. While the non-migratory population is ecologically unrealistic, its output provides a yearly baseline for foraging patterns and krill intake over the modeled foraging season. Transition probabilities between *S*_*1*_ and *S*_*2*_ are identical across all presented models, thus the null non-migratory agents receive the same foraging and movement cues as the southward migration models. A yearday-driven migration strategy was also tested as a null model, and results can be found in the supplement.

### Population Comparison Metrics

Population-level behaviors of each model are compared against empirical data of migration timings and the null model. First, the modeled southward migrations are compared against empirical data from the Monterey Bay hydrophone. Foraging season durations are known to be latitude dependent (Calambokidis *et al*. 2015), hence hydrophone recordings are compared to the subset of modeled agents whose migration initiated north of Monterey Bay (north of 36° N). Modelled migration mechanisms are characterized as realistic if the resultant median southward migration dates align with empirical measurements in terms of the correct date range (Oct-Nov) and an interannual flexibility on the order of months. A Mann-Whitney U-test (Mann & Whitney 1947) is applied with a null hypothesis that the median migration dates for a modeled strategy have the same distribution as the hydrophone dates. The alternative hypothesis that the model migration dates are from a distribution with earlier migration times.

An agent’s total annual krill intake is defined as the sum of the krill densities at all foraging locations. As a second metric, we compare a population’s total krill intake against that of the hypothetical non-migratory population (*m*_0_). Prey levels are dynamic across years and *m*_0_ acts as a null model representing the maximum yearly krill intake the population could achieve if agents did not migrate and made full use of all the available resources. To compare results across years, yearly krill intakes for each population are represented as a percentage of the median intake of *m*_0_. While the total krill intake provides a useful measure of energy intake and foraging success, this metric is maximized by delayed migrations.

To understand the flexibility of migration strategies, performance metrics are analyzed when separated by yearly environmental conditions. The median total krill intake of *m*_1_ is used to categorize each year as low, average, or high krill availability (Supplementary Table 1).

### Parameter fitting, sensitivity analysis, model robustness

All simulations were initialized with 2,000 agents to mimic the population of Northeastern Pacific blue whales. IBM outputs are naturally stochastic and each simulation, even with a fixed parameter set, represents one possible model outcome or realization. To account for the inherent stochasticity and ensure that results are representative of the full range of possibilities, at least 100 simulations were run for each year in 1990-2010 and outcomes report the aggregation of all simulations.

Latin Hypercube Sampling (Marino *et al*. 2008) was conducted to test the sensitivity of model outcomes to parameter values, in particular the sensitivity of southward migration statistics and distributions. The sensitivity analysis aided in selection of model parameters; details are in the Supplement.

## RESULTS

Southward migration decisions based solely on individual cues (personal information; *m*_per_) yield unrealistically early migrations (Figure 2). Median migration dates of *m*_per_ span yearday 209 (late-July) through yearday 283 (mid-October) and are significantly different from those of the empirical dataset (Mann-Whitney U-test p-value<0.001; Supplementary Table 5).

**Figure 2:**
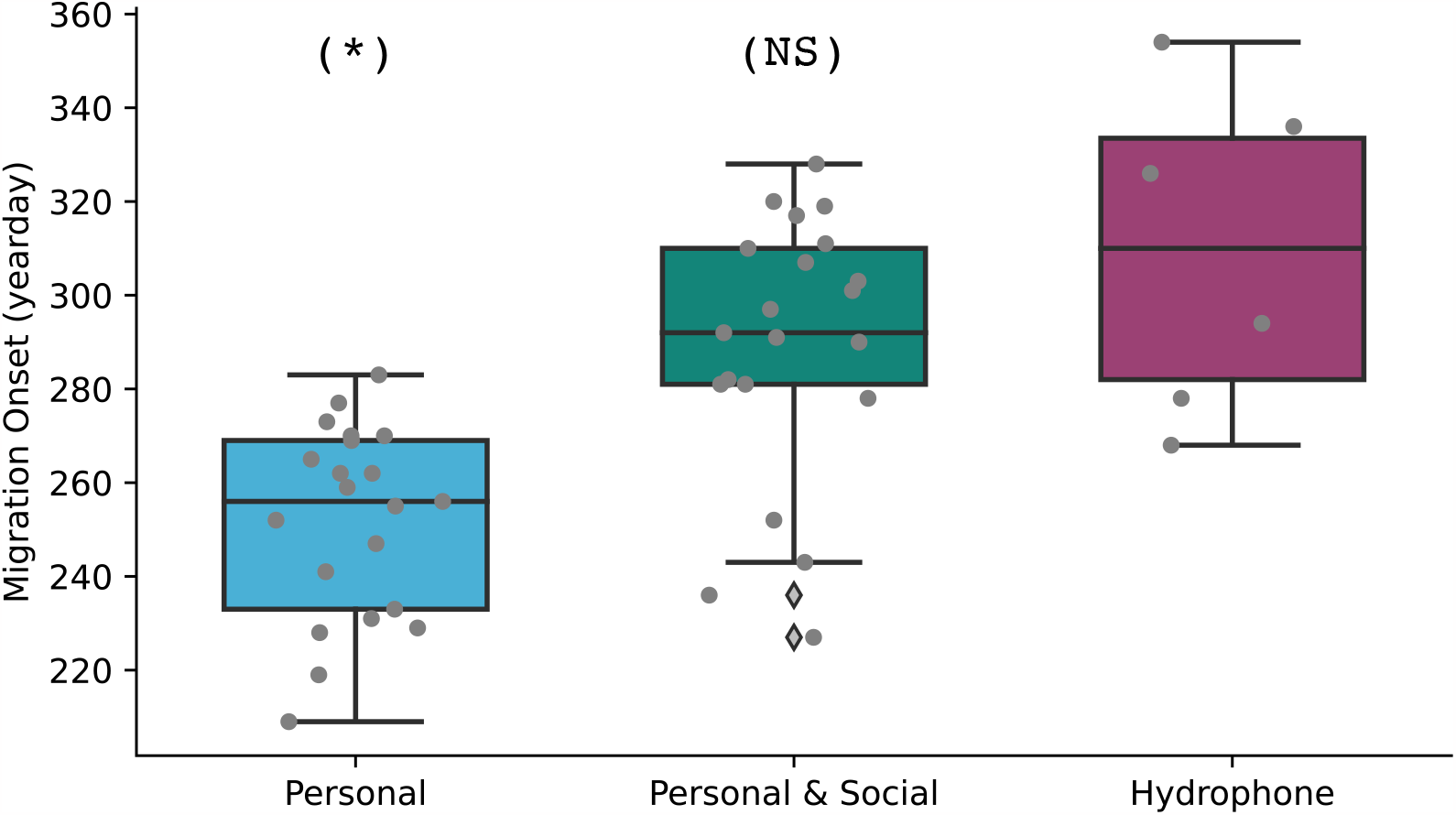
Median migration dates from Monterey Bay hydrophone data and modeled migration mechanisms. Migration statistics for each modeled migration mechanism were calculated using the subset of the agents whose migration initiated north of Monterey Bay. Boxplots show the distribution of annual median migration dates (shown in gray dots). The (*) label indicates a statistically significant difference between the set of median migration dates of the modeled mechanism and the hydrophone dataset and the (NS) label indicates no significant difference.

Realistic, late-season migrations that overlap with empirically-observed migration timing emerge when social communication is added to the decision process. For the 21 simulated years, the median migration dates of *m*_per&soc_ span late August (yearday 236) through early November (yearday 319).

Median migration dates of *m*_per&soc_ fall after yearday 281 (early October) in 75% of the simulated years and are statistically indistinguishable from the timing of southward migration empirically observed from the Monterey Bay hydrophone (p-values=0.16; Figure 2; Supplementary Table 5). Further, a sensitivity analysis revealed that trends in migration timings are robust across a wide range of model parameters (Supplementary Figure 1).

The relative krill intake of each modeled population is compared to the null non-migratory simulations (*m*_0_) in Figure 3 with years separated by prey availability. Across all prey conditions, populations using social information (*m*_per&soc_) consume a greater total amount of krill than those using the asocial strategy (*m*_per_). In years with low krill availability, *m*_per_ consumes only 49% of the *m*_0_ krill intake compared to the *m*_per&soc_ median of 69.8% of the available krill. These trends continue into the average and high krill years, with the asocial population consuming a median of 57.8% and 73.5% of the *m*_0_ intake, with the socially informed population consuming a median of 85–92% (Figure 3, Supplementary Figure 5).

**Figure 3:**
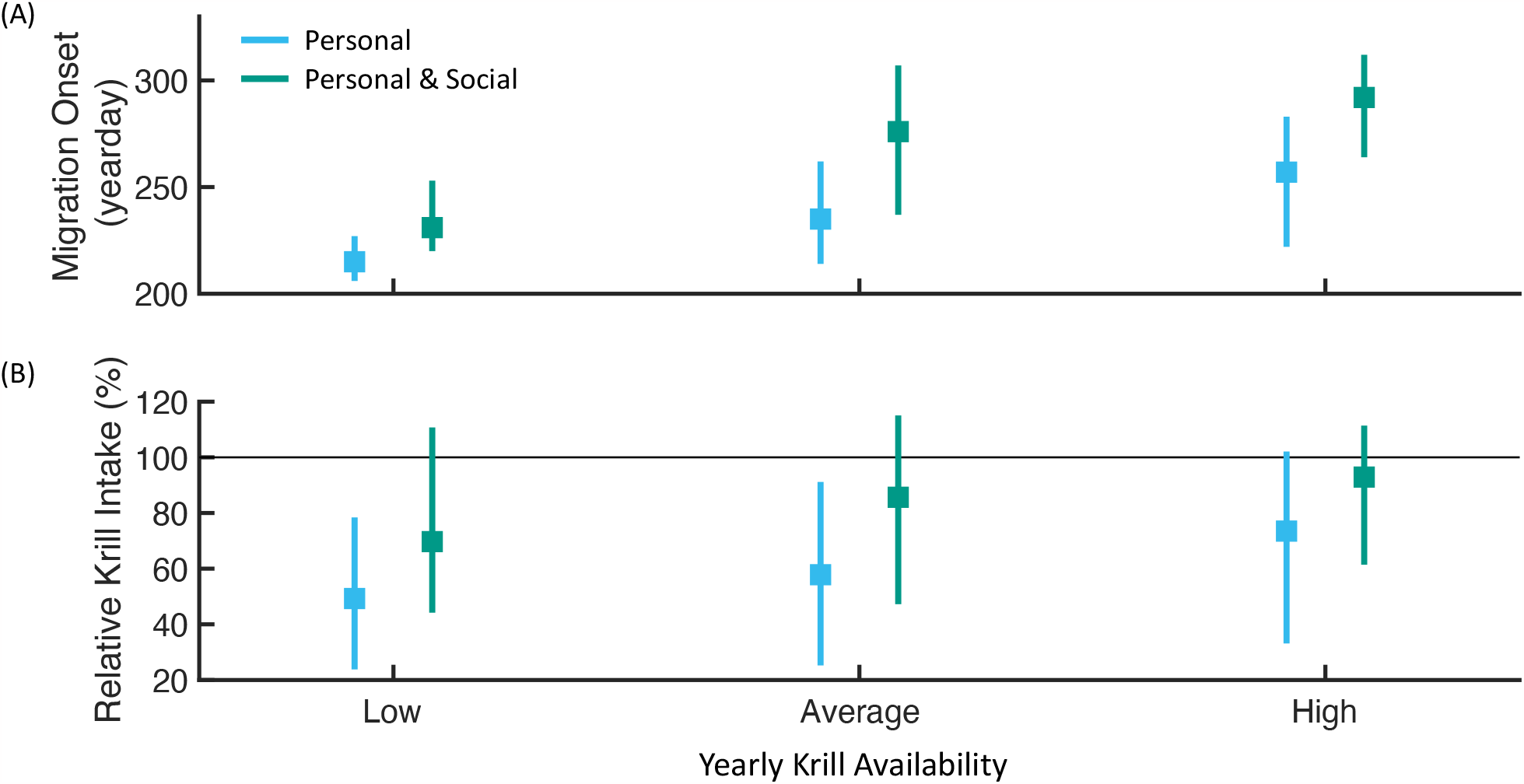
Southward migration distributions and krill intake for modeled populations 498 separated by krill availability (low, average, high). Boxplots show (A) IQR of migration distributions and (B) relative krill intake for each migration mechanism. Values in (B) are computed as a percentage of the total non-migratory (null) population intake. Grey line indicates the median intake of the null population. Results are aggregated by yearly krill availability on the x-axis.

The importance of the social communication range was tested by varying the maximum call radius for a range of detection distances (0-500 km, Figure 4). A maximum call radius of 0 km corresponds to the *m*_per_ strategy. The timing of southward migrations converges to a late season median as the maximum call radius increases. Increasing call radius further leads to a decrease in population-wide variation in migration timing, as indicated by the narrowing of the migration timing distribution. Likewise, the increase in call radius leads to a krill intake nearly 175% of the personal-only strategy (Figure 4B). A relatively small maximum call radius of 5 km leads to a median 12-day delay in departure date and median prey consumption nearly 22% greater than the *m*_per_ krill intake. Maximum call radii of 25 km and above yield at least 50% greater prey intake than *m*_per_ and median migration dates a full 30 days later.

**Figure 4:**
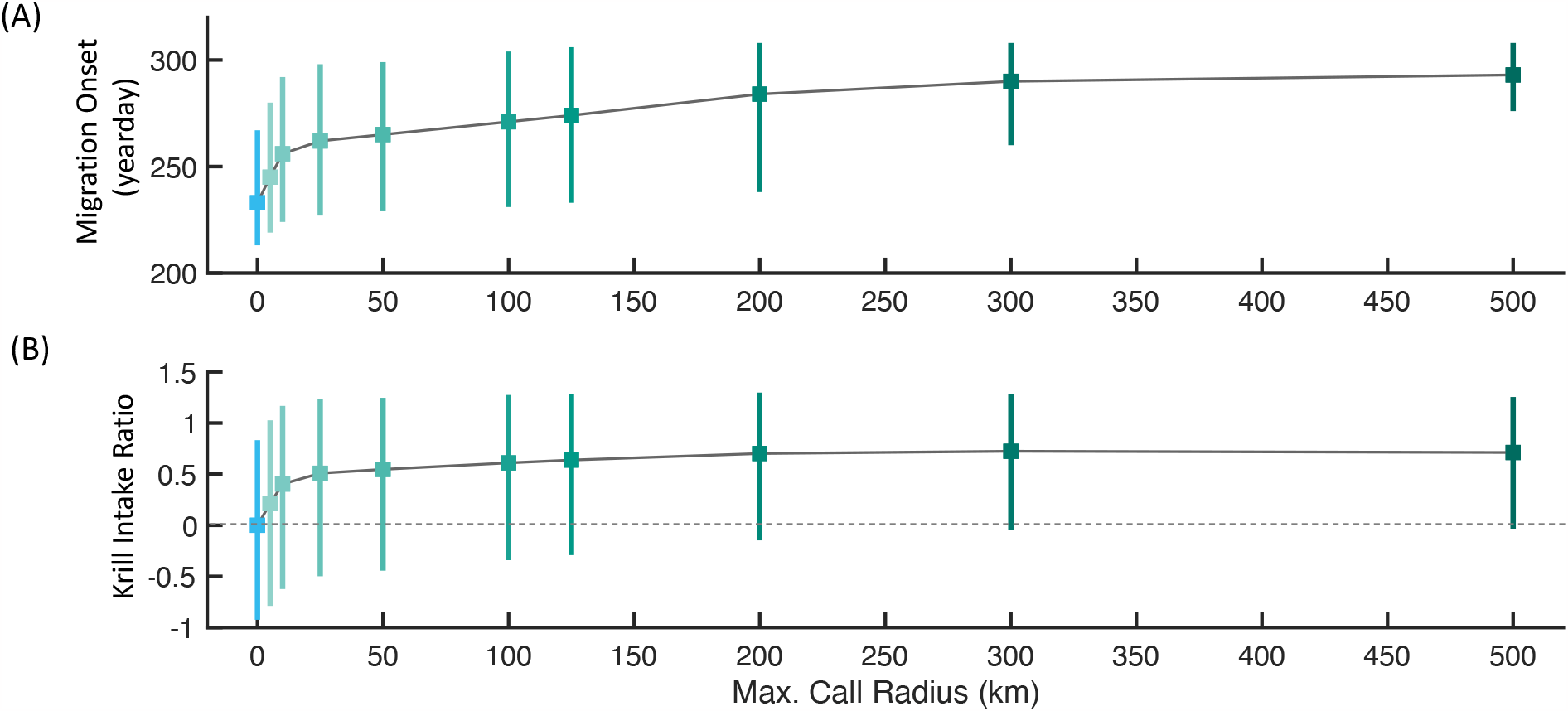
Impact of maximum call radius on migration timing and krill intake. Boxplots show (A) IQR of migration distributions and (B) relative krill intake compared to the personal-only population. Values in (B) are computed as a ratio of the deviation from the total personal population m_per_ intake. Data aggregated across all 21-years of ROMS data. A maximum call radius of 0 km corresponds to the m_per_ population.

## DISCUSSION

Understanding the mechanisms of migration is a longstanding question in ecology (Dittman & Quinn 1996; Lohmann *et al*. 2001; Flack *et al*. 2018). A growing body of research has focused on the role of social interactions in migratory decisions (Aikens *et al*. 2022; Oestreich *et al*. 2022a; Jesmer *et al*. 2018; Mueller *et al*. 2013), as they can enable populations to navigate noisy environmental gradients and can give rise to collective migrations (Guttal & Couzin 2010). Yet whether such social cues enable collective migrations and associated fitness benefits at the extreme spatial scale of dynamic pelagic ecosystems has remained untested. Our results indicate that long-distance social interactions are likely an important ingredient in driving collective migration in a long-distance ocean migrant, enabling flexible migration timing and associated benefits in foraging performance.

The recent discovery of an acoustic signature of blue whales’ transition from foraging to migration (Oestreich *et al*. 2020) prompts the hypotheses that (1) blue whales incorporate long-distance social information from conspecifics to time their migratory departure from foraging habitat; and (2) this social information improves population-wide foraging outcomes. Using a series of IBMs in conjunction with empirical observations, we find support for these hypotheses, showing that only models including a socially-informed strategy enable flexible and realistic migratory departure timing from blue whales’ vast and dynamic foraging habitat (Figure 2) across a broad range of realistic model parameter values (Supplement). This use of long-distance information from conspecifics enhances individuals’ foraging performance under interannual ecosystem variation (Figure 3).

Optimal foraging theory posits that animals will optimally choose, allocate time to, and depart resource patches in ways that maximize individual fitness (Pyke *et al*. 1977, Charnov *et al*. 1976). However, this model assumes that animals have complete, landscape-wide knowledge of resource availability – an assumption that is typically violated in nature and is influenced by both patch predictability and individual sensing capabilities. Our results are consistent with theory that predicts the value of nonlocal information to be high in systems with highly-transient resource patches, enabling more optimal foraging decisions despite high environmental unpredictability (Fagan *et al*. 2017). Many pelagic ecosystems –including blue whales’ foraging habitat in the Northeast Pacific (Santora *et al*. 2011) – display these resource dynamics (Marquet *et al*. 1993; Benoit-Bird & McManus 2012; Hazen *et al*. 2013; Bertrand *et al*. 2014), suggesting that nonlocal information should be particularly valuable in this setting. Moreover, our study indicates that theoretical predictions about the utility of social information in heterogenous ecosystems extend to the extreme spatial scale and dynamism of the pelagic ocean. Modeled individuals considering only personal information depart when they experience poor forage conditions locally, even if high-quality foraging opportunities remain elsewhere. This lack of awareness of broader forage availability leads asocial individuals to migrate earlier than empirical observations show (Figure 2) and results in poorer foraging performance relative to social migrants (Figure 3B). Delayed migration is enabled by an acoustic social network informing the population about broader forage availability than can be sensed individually (Figure 3B), thus enabling more optimal foraging decisions. The shortcomings of asocial strategies and success of socially-informed migratory decisions are robust to model variation and parameterization (Supplementary Figure 3). Hence, models including socially-informed migratory decisions can consistently better explain trends seen in empirical data relative to asocial-only models, suggesting that long-distance social information is an important and necessary source of information in blue whales’ decision of when to depart from foraging habitat and initiate breeding migration.

Social information enhances the range of nonlocal information, but the information acquired personally by information-sharing individuals is often highly-correlated (Winklmayr *et al*. 2023). Such homogeneous personal information among individuals can limit the value of social information and the ability of groups to collectively sense broad-scale ecosystem gradients or changes (Kao & Couzin 2014). In pelagic ecosystems where ephemeral resource patches display heterogeneity across a broad range of spatial scales (Marquet *et al*. 1993; Hazen *et al*. 2013), effective collective sensing of broad-scale ecosystem dynamics therefore would require long-distance information transfer between individuals. Blue whales have evolved to take advantage of the potential for long-distance acoustic communication in aqueous medium of pelagic ecosystems (Au & Hastings 2008), producing extremely loud and low frequency songs which are detectable over at least tens (if not hundreds) of kilometers (Oestreich *et al*. 2020). These physical attributes of blue whales’ songs allow for the long-distance communication necessary for collective sensing and migration in the open ocean. While the precise maximum communication range between blue whales is not known, implementing even relatively conservative estimates results in realistically-delayed migration timing and enhanced foraging performance relative to a hypothetical asocial population (Figure 4). Even as the social communication radius grows and population-wide migration timing becomes increasingly collective (decreasing IQR in Figure 4A), the marginal value of social information from increasingly-distant conspecifics for foraging performance approaches an asymptote (Figure 4B). This pattern likely arises from information saturation, where increasingly distant calls provide limited additional information about foraging conditions. As a result, individuals accrue the majority of foraging benefits from social information transfer at moderate ranges of vocal communication (< 50 km; Figure 4) relative to the detection range of these signals above background noise (at least from human listening devices; Oestreich *et al*. 2020).

These findings also highlight how the sources of information driving migratory decisions vary depending on the relative value of distinct information sources during different phases of an annual migratory cycle. As Northeast Pacific blue whales migrate northward in the early spring, individuals have limited personal information about contemporaneous forage conditions, leading to a reliance on spatial memory of long-term foraging hotspots (Abrahms *et al*. 2019). Moreover, social information transfer via song is sparse during the early foraging season with individuals producing little song during the summer months (Širović *et al*. 2015; Oestreich *et al*. 2020; Pearson *et al*. 2023). As the foraging season progresses and krill prey become less abundant (Croll *et al*. 2005; Fossette *et al*. 2017), the relative value of long-distance social information should increase as individuals must determine if local prey depletion is representative of broader forage availability. Indeed, song production increases throughout the foraging season, reaching a peak in the late fall when this population begins southward migration on average (Oestreich *et al*. 2020). While this social information transfer provides clear benefits to receivers (Figure 3), the benefits to signalers are not immediately clear. In this situation, one might expect “cheating” to evolve unless song production provides some other fitness benefit to signalers. Song in baleen whales is widely associated with reproduction as it is believed to be a male-specific behavior and peaks immediately preceding the breeding season (Oleson *et al*. 2007). Even if the primary purpose of song relates to reproduction, the diel patterns of blue whale song create secondary information on behavioral state due to diel partitioning of feeding and singing behavior during the foraging state (Oestreich *et al*. 2020). In this way, the information on behavioral state encoded in diel patterns of blue whale song provides unintentional social information about forage availability to eavesdropping conspecifics. The use of such public information is widespread in animals’ behavioral decisions (Valone & Templeton 2002; Danchin *et al*. 2004), and its inadvertent nature allows for evolutionary stability even without benefit to the signaler (Dall *et al*. 2005).

Elucidating how animals use various information sources in their migrations advances our understanding of movement ecology in theory and in practice. Discovery of blue whales’ probable use of long-distance social information improves our understanding of this endangered population’s ability to adapt to changing ecosystem conditions in the Anthropocene. Blue whales’ reliance on memory during the northward migration onto foraging grounds suggests limited capacity for this population to adapt to rapid environmental change in this portion of their annual cycle (Abrahms *et al*. 2019). Yet the ability to flexibly track interannual variation in ecosystem phenology when departing on migration from foraging grounds suggests greater ability to adapt to environmental change. Our findings suggest that this adaptive flexibility is underpinned by long-distance communication—as a result, the already-detrimental impacts of noise pollution on this population might also impair the use of social information in migration timing and its associated foraging benefits (Duarte et al. 2021). Growing calls and proposed solutions for reducing human impacts on marine soundscapes (Duarte, et al. 2021; ZoBell *et al*. 2021, 2023; Findlay, et al. 2023) hold promise for mitigating this impact, aiding in the conservation of marine megafauna and collective migrations. Finally, our results point to a growing need to understand the function and consequences of long-range social communication in migratory populations navigating rapidly-changing ecosystems.

## Supplementary Information

## 1 Overview, Design concepts, and Details (ODD)

This section of the supplementary information is formatted under the Overview, Design concepts, and Details (ODD) protocol (Grimm et al., 2006; Grimm et al., 2010; Grimm et al., 2020).

### 1.1 Purpose

The purpose of this individual based model (IBM) is to investigate the role of foraging and long-range social information in driving the yearly southward breeding migration of Northeastern Pacific blue whales.

### 1.2 Entities, state variables, and scales

Entities or agents in the IBM represent a single whale. State-variables track how each agent moves through and experiences the domain. The domain spans 116-128^*°*^W and 32-44^*°*^N and is divided into 3 km × 3 km spatial patches. Each patch is assigned a sea surface temperature (SST; ^*°*^C) and near surface krill abundance. The SST and krill density are updated daily (24 hours) and are provided from an implementation of ROMS (see 1.6).

To investigate drivers of southward migration, we use a series of IBMs, each of which is designed with a distinct southward migration mechanism based on a combination of environmental and social information. The IBMs are formulated as a state-switching model with four behavioral states representing transit and forage behaviors during the foraging season and breeding migration. In all models, each agent is assigned the following state-variables: behavioral state, location, SST, and krill (Table 1). In models with social calls the three additional state variables of sex, calling behavior, and received call signals are assigned. One time step represents 6 hours (4 time steps/day) and simulations were run for 180 days. Each simulations includes 2,000 agents. Simulations are run for years 1990-2010.

**Table 1:**
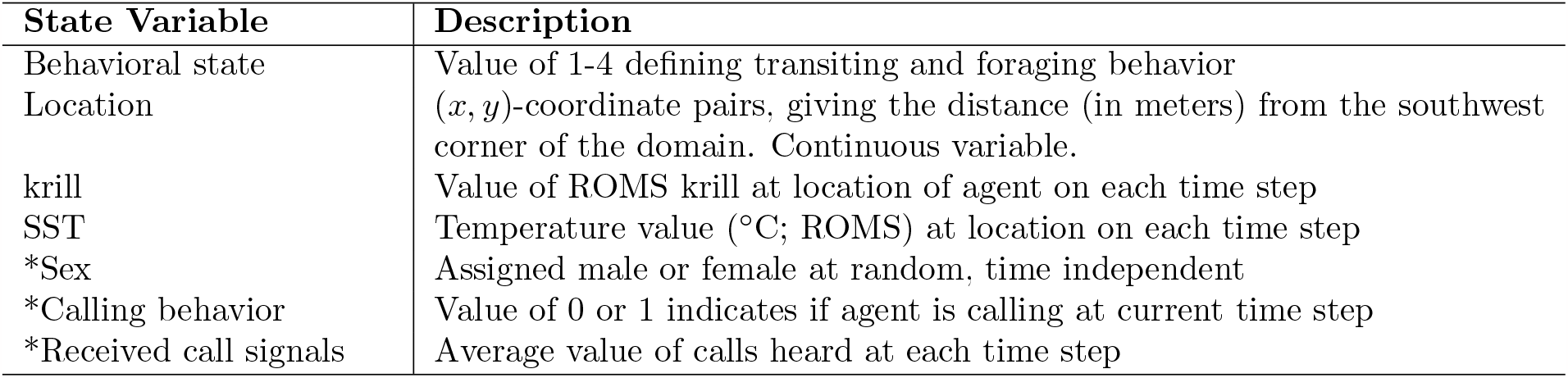
Description of state-variables defined to each agent. Variables are unitless, unless otherwise specified. An asterisk next to the state variable name indicates a variable that is only used in models with social calls.

### 1.3 Process overview and scheduling

The model progresses in 6 hour time steps. Within each time step, the state-variables for each agent are updated following the order displayed in Figure 1. Environmental state variables are updated every 24 hours (4 time steps). Details of each process are included in Section 2. Movement updates are selected from state-dependent step length and turning angle distributions.

**Figure 1.**
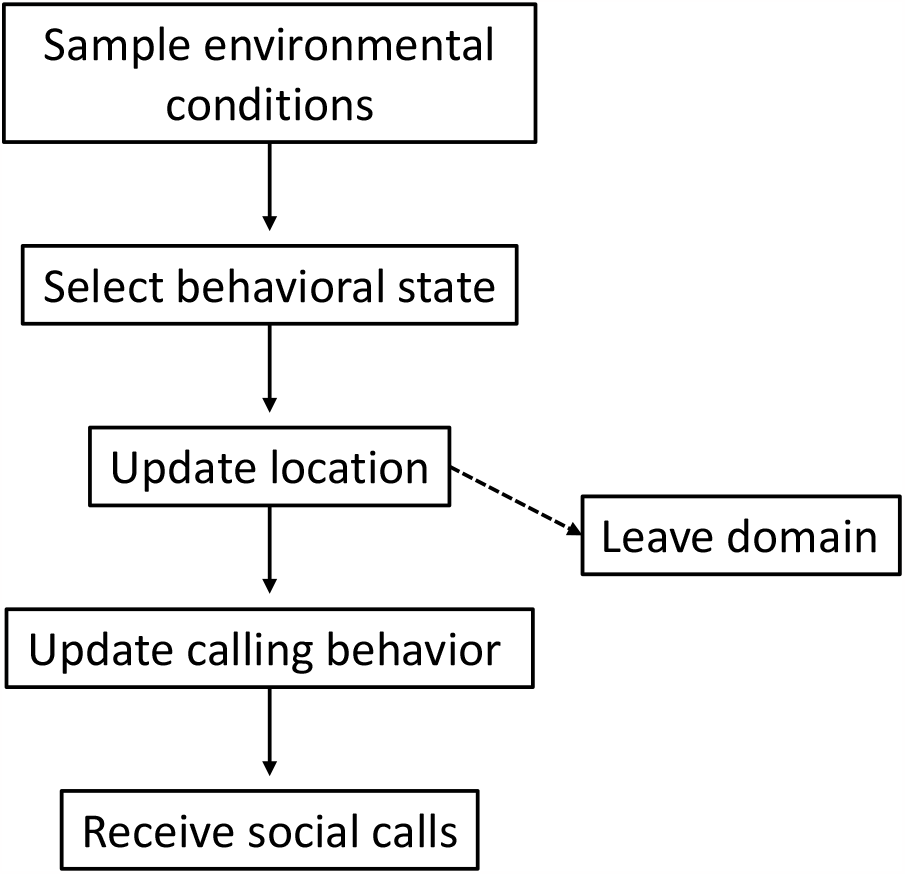
Schematic of underlying IBM algorithm. Process represents a single time step and is completed for each agent.

### 1.4 Design concepts

#### Basic Principles

The presented IBM takes the form of a state-switching model, with behavioral states capturing the distinct movements associated with transiting, foraging, and migrating behaviors. Behavioral states and their associated movement distributions are based on tagging data (Bailey et al., 2009) and have been successfully incorporated into IBMs (Dodson et al., 2020).

In this study, multiple southward migration mechanisms based on environmental and social cues are tested via the state transition functions that control how agents enter the migratory states. Migration mechanisms are evaluated by comparing the timing of modeled southward migrations to empirical hydrophone observations (Oestreich et al., 2022; Oestreich et al., 2020). Thus, the model will provide insights about the processes underlying blue whale migrations.

#### Emergence

The existence and timing of the southward breeding migration is an emergent behavior that arises from foraging behaviors and received social calls. The emergent southward migration is the main output of interest. Additional emergent population-level behaviors include seasonal foraging trends and locations.

#### Adaptation

Agents update their behavioral state on each time step based on the environmental conditions and received social information. Behavioral states dictate the step length and turning angle of the next movement update, mimicking movements associated with transit and forage behaviors (Bailey et al., 2009).

#### Objectives

There are no model processes that seek to maximize some objective.

#### Learning

Agents do not learn throughout the model implementation.

#### Prediction

Agents do not make any predictions of future consequences of their decisions.

#### Sensing

It is assumed that agents can locally sense the environmental state variables. Agents are able to sense the SST and krill values of neighboring grid cells and can determine the direction of highest foraging probability (if one exists).

Agents can sense and receive social calls within a maximum radius of their location. The default radius is set at 125 km. Agents are assumed to be able to sense ocean versus land. Movement updates that bring agents onto land are not allowed.

#### Interaction

Agents interact through social calls. Calls include information about the call sender’s recent foraging behaviors (see Section 2). The information shared impacts southward migration decisions. In models without social communication, agents do not interact.

#### Stochasticity

The selection of behavioral states and movement updates is inherently stochastic. Behavioral states are selected probabilistically given a state transition matrix. Movement updates are selected randomly from associated turning angle and step length distributions. Stochasticity is used to reproduce variability in the modeled decisions and outcomes.

#### Collectives

Agents do not belong to any collective unit.

#### Observation

During model simulations, the behavioral state, cumulative krill intake, average recent krill intake, calling behavior, and received calls are recorded for each agent on each time step. These quantities contribute to the selection of behavioral states and provide a useful comparison between distinct migration mechanisms.

Calling behavior only impacts the behavioral states and decisions of agents in the models with social behavior. However, in practice, calling behavior can be monitored in all models.

### 1.5 Initialization

Since we are interested in factors driving the breeding migration, models are initiated on July 1st. Agents’ initial locations are selected uniformly at random within areas of climatologically high krill densities from the ROMS data Abrahms et al., 2019. Regions of high climatologically krill were defined by averaging the krill abundance on July 1st 1990-2010 for each spatial grid cell, then thresholding the averages.

At the start, 50% of agents are randomly assigned to be male, an important fact since the primary producer of long-distance song are males.

### 1.6 Input data

The environmental variables SST and near-surface krill abundance are precomputed and supplied as inputdata to the IBM. These variables are precomputed using a Regional Ocean Modeling System (ROMS) implementation that is specifically parameterized for the California Current region (Fiechter et al. 2018; Fiechter et al. 2020). Additionally, the ROMS implementation has been coupled with NEMUCSC, a biogeochemical model adapted from the North Pacific Ecosystem Model for Understanding Regional Oceanography of Kishi et al. (2007), and provides the krill concentrations. ROMS-NEMUCSC data that spans the domain is available for the years 1990-2010 at 3km spatial resolution and daily temporal resolution. Simulated krill concentrations have been evaluated against existing in situ data for May-June and are expected to adequately reproduce observed krill aggregation regions during the upwelling season (Fiechter et al., 2020). This data is available at https://doi.org/10.7291/D1KD4J.

## 2 Submodels and Model Details

This section represents the *Submodels* portion of the ODD protocol. Subsections define the processes in Figure 1.

Throughout the model description, the subscript *n* will indicate a quantity or variable for an individual agent, where *n* ∈ {1, 2, …, 2000}. Additionally, *t* is used to represent the time step of the simulation. Model time steps are 6-hours in length and simulations are initiated on July 1 (*t*_0_). The yearday *t*^∼^ can be extracted from the time step *t* using the floor-function as 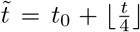. Variables are summarized in Table 2 and notation is consistent with that of the main text.

**Table 2:**
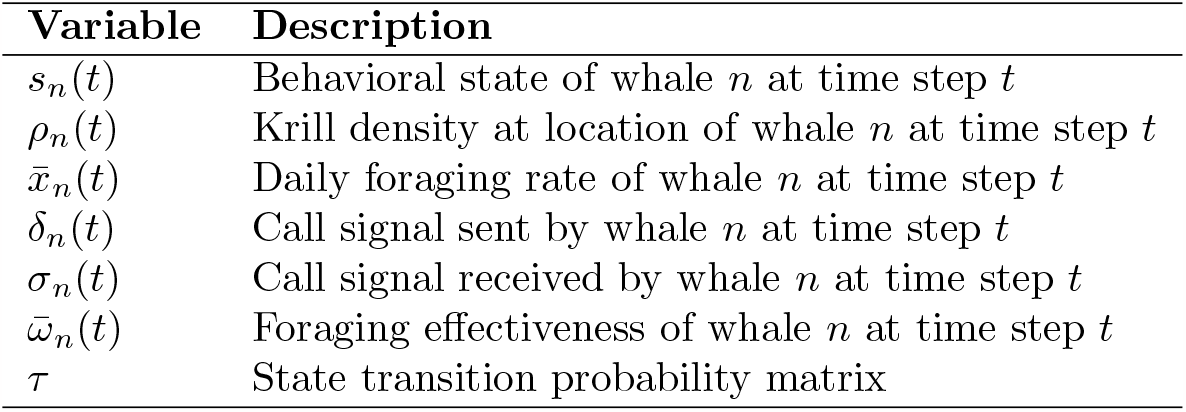
Model variables. Summary and descriptions of model variables.

### 2.1 Social Communication

The social communication process is divided into the call producers and call receivers.

#### 2.1.1 Call Producers

The call signal sent by agent *n* on at time step *t* is defined as

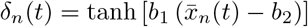

where 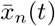 denotes the average 24-hour foraging behavior

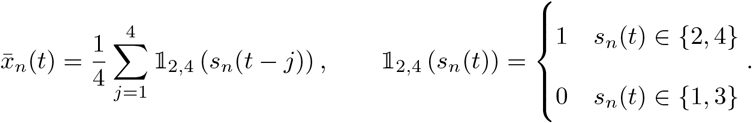

Here, 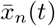 takes on a value between 0 and 1, with 0 representing all transit and 1 all forage behaviors. The call signal *δ*_*n*_(*t*) is intended to be a time-dependent measure of foraging success analogous to the ratio CI_night_:CI_day_ (Oestreich et al., 2022; Oestreich et al., 2020).

At the start of the simulation, 50% of agents are randomly assigned as male (call producers) and on each time step, a subset of male whales are randomly selected to produce calls. The proportion of male whales calling on each time step increases linearly from 5% on July 1 to 30% on December 31.

#### 2.1.2 Call Receivers

Define *M*^*n*^(*t*) to be the subset of whales whose calls are heard by whale *n* at *t, N*_*n*_(*t*) = |*M*_*n*_(*t*)| (cardinality of the set *M*_*n*_(*t*)) to be the number of calls received, and *R*_*j,n*_(*t*) the distance (in meters) between two agents *n* and *j*. Then, the call signal received by whale *n* at time *t* is an average of the all calls received, weighted by the inverse-square law dependent decay of the call amplitude

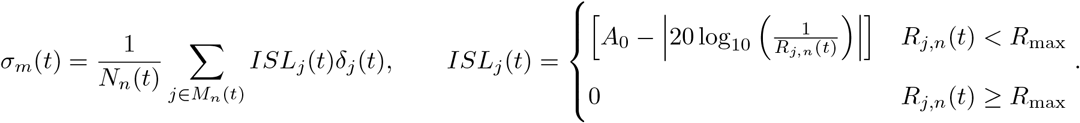

Call signals are capped at a maximum radius *R*_max_. Calling behaviors and the inverse-square law amplitude decay are independent of call radii.

### 2.2 Selecting behavioral states

The four behavioral states S_1,2,3,4_ represent transiting and foraging behaviors during the northward foraging and southward breeding migrations. Each behavioral state is associated with characteristic movements defined by step length and turning angle distributions.

Transitions between the four behavioral states S_1,2,3,4_ are governed by the state transition probability matrix

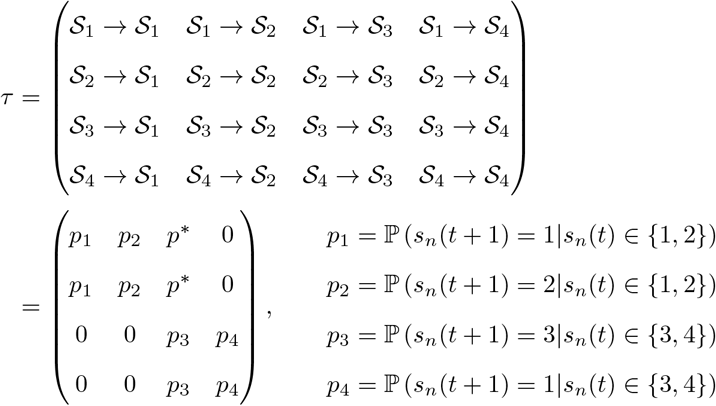

where *s*_*n*_(*t*) ∈ {1, 2, 3, 4} is the behavioral state of whale *n* at time step *t*. Allowed transitions are show in the schematic in Figure 1 in the main text.

Southward migration strategies are incorporated in the southward transition probability *p*^*∗*^. Tested migration strategies are based on a combination of an agent’s personal foraging behavior and received social information. Additionally, let 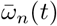 be the average foraging behavior and 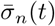 be the average received social information of agent *n* over a period of *T* = 40 time steps (10 days). These are defined by

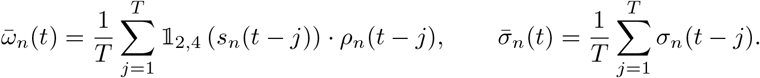

Migration strategies are encoded in the transition probability *p*^*∗*^ = P (*s*_*n*_(*t* + 1) = 3|*s*_*n*_(*t*) ∈ {1, 2}). The transition probability functions associated with each strategy are given below. Although the subscript *n* is omitted, all transition probabilities are assigned for each agent. All parameters are included in Table 3.

**Table 3:**
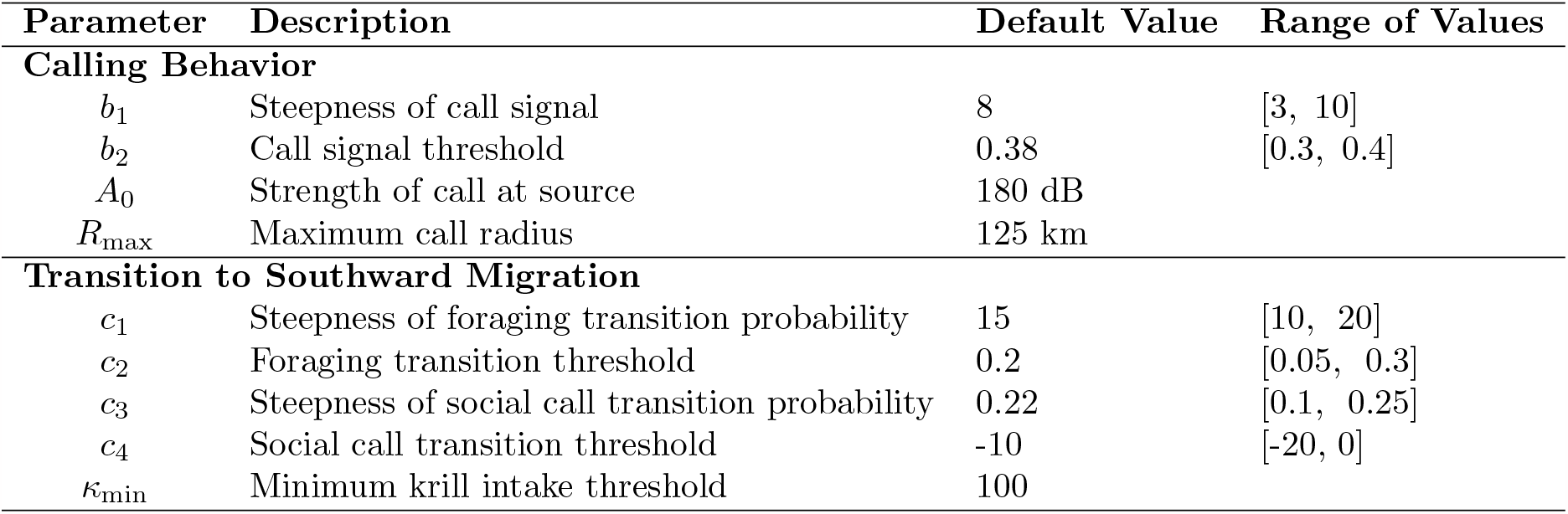
Calling and migration model parameters. Summary and descriptions of model parameters related to calling behavior and southward migration. Model results computed using default parameter values. Sensitivity analysis conducted over the listed range of values. Parameter values with no listed range were not included as part of the sensitivity analysis. Parameters without units are unitless.

1. Individual foraging efficiency (personal):

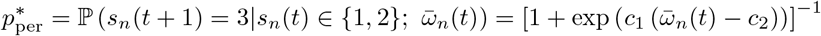
2. Individual foraging efficiency and minimum krill intake (personal & min krill):

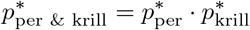
3. Social communication (social):

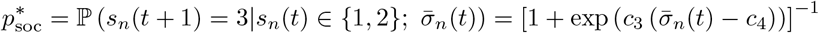
4. Individual foraging efficiency and social communication (personal & social):

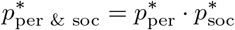

The personal, social, and personal & social strategies and their results are described in detail in the main text. In the Supplementary Results (Section 8), we additionally include results of a migration strategy with a minimum krill intake requirement. Specifically, agents are only able to migrate if they exceed a minimum krill threshold *κ*_min_. Thus, define the function

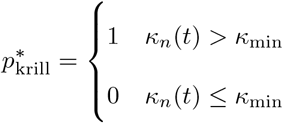

where the cumulative krill intake *κ*_*n*_(*t*) is found by summing the krill density *ρ* at the agent’s foraging locations

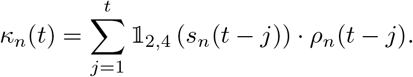

State transition probabilities are computed as follows. First, *p*^*∗*^ is computed and fixed. Then, for agents in S_1,2_ the probabilities *p*_1_ and *p*_2_ are defined using the fact that the rows of the STPM sum to 1. Thus, the probabilities are set to

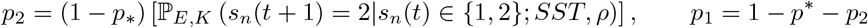

where P_*E,K*_ (*s*_*n*_(*t* + 1) = 2|*s*_*n*_(*t*) ∈ {1, 2}; *SST, ρ*) is the probability of foraging due to SST and krill density defined in (Dodson et al., 2020). Parameter values for the forage-transit selection process are identical to those in (Dodson et al., 2020). Transition probability functions between S_1_ and S_2_ are identical across all presented models.

For agents in S_3,4_, we likewise define the probability of foraging and utilize that *p*_3_ + *p*_4_ = 1. Thus,

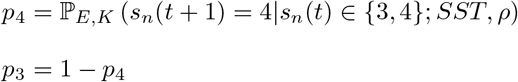

where *p*_4_ has a stricter foraging threshold (high krill density required for foraging). Transition probabilities between S_3_ and S_4_ are identical across all presented models that include the southward behavioral states.

#### 2.2.1 Sampling Environmental Conditions and Location Updates

On each time step, agents sample and store as state variables the SST and krill density at their current location (values from ROMS data). These environmental parameters are used to compute the elements of the state transition probability matrix.

Regardless of migration strategy, movement updates are selected from distinct turning angle and step length distributions associated with characteristic transit and forage behaviors. See Dodson et al., 2020 for movement distributions.

## 3 Model Assumptions

Here, we summarize and clarify model assumptions.

- Behavioral states and positions are updated every six hours. Transition probabilities within the north and southward migration categories (between states S_1_ → S_2_ and S_3_ → S_4_) are based only on SST and prey levels.
- Transition probabilities between S_1_ → S_2_ are identical across all presented models.
- Transition probabilities between S_3_ → S_4_ are identical across all presented models that include the southward behavioral states.
- Individuals commit to southward migration and are not permitted to transition from states S_3,4_ to S_1,2_.
- Agents are not bound to the domain and will freely leave if their movement updates take them outside the domain.
- Since we are interested in factors driving the breeding migration, the model is initiated on July 1st. Agents’ initial locations are selected uniformly at random within areas of climatologically high krill densities from the ROMS data (Abrahms et al., 2019).
- Tested migration mechanisms represent strategies based only on foraging history, information gathered from social calls, and year-day (null model).

## 4 Null Models

Model results are tested against two null models. The first null model, a hypothetical non-migratory population, is described in the main text and is the two-state model of Dodson et al., 2020. The second null model follows a yearday-driven migration strategy. Migration dates for individuals in the yearday model are pre-determined and follow a normal distribution with a mean of 310 (November 6) and standard deviation of 20 days. The mean and standard deviation represent average historical trends in blue whale migrations and average call behavior recorded from the MARS hydrophone (Oestreich et al., 2022; Oestreich et al., 2020).

## 5 Software

All simulations of the IBM were programmed in MATLAB and run using MATLAB Version 9.13 (R2022b) (The MathWorks Inc., 2022a). Since each simulation run is independent from others, the simulations were run in parallel using the the parallel computing toolbox (The MathWorks Inc., 2022b). Code for simulating and processing the individual-based model is available at https://doi.org/10.5281/zenodo.8305222.

Statistical analysis using the Mann-Whitney U-test (Mann and Whitney, 1947) was conducted with Python Version 3.10.8 using the mannwhitneyu function from the stats module of SciPy (Virtanen et al., 2020).

## 6 Yearly Krill Availability

The 1990-2010 ROMS-NEMUCSC data was separated into years of low, average, and high krill availability based on the median total krill intake of the null non-migratory population. Years were classified as “average” if the median krill intake fell roughly in the middle 50% of the data. Yearly classifications of krill availability are included in Table 4 and ranked median total krill intakes are in Figure 2.

**Table 4:**
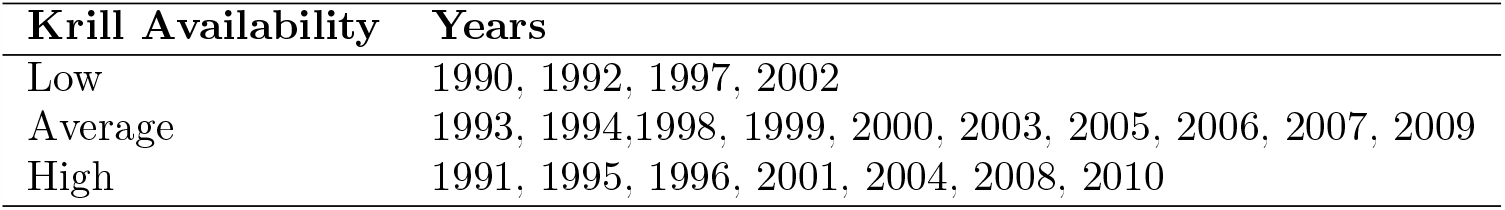
Classification of years based on krill intake of the null non-migratory population.

**Figure 2.**
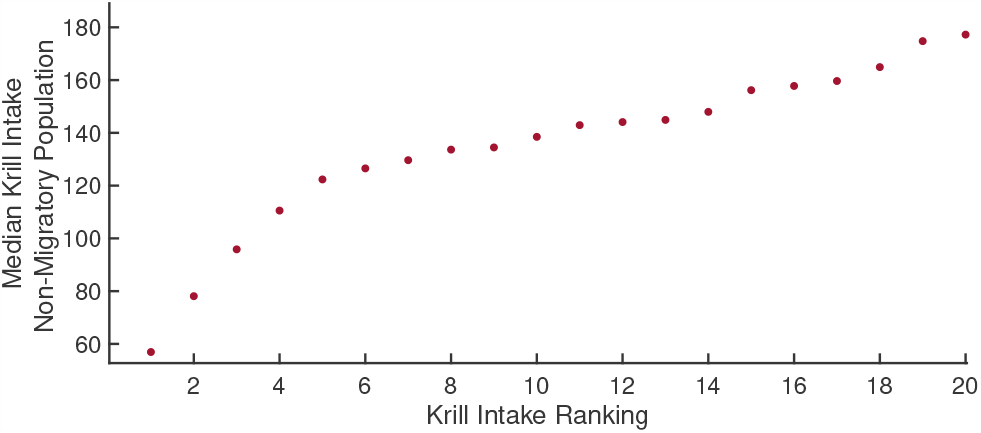
Yearly krill intake. Ranked median yearly krill intake of the null non-migratory population.

## 7 Sensitivity analysis

Robustness of the timing of the migration mechanism was tested with using random parameter samples. Realistic, but large intervals were set for all parameters (Table 3). A total of 1,000 trials were run for each year in 2000-2010 with parameters randomly selected from the set parameter ranges using Latin Hypercube Sampling procedures (Marino et al., 2008). Figure 3 shows the median migration dates for all random sample trials. Tested parameter intervals were consistent across all strategies.

The addition of social calls leads to late season migrations over the full range of realistic parameters. Notably, even the lowest median migrate dates for social strategies are within reason. The asocial strategies consistently lead to early migrations over the broad range of parameters with median dates as early as September-even when restricted to the subset of the population that migrates from the higher latitudes. The results of this sensitivity analysis are important - they show that including social information in migration decisions leads to a late season migration over a broad range of parameters and conditions.

The default set of parameters stated in Table 3 and used for the results in the main text were selected to give later season migrations in the personal-only population (migration timings in the upper tails of the distributions in Figure 3).

**Figure 3.**
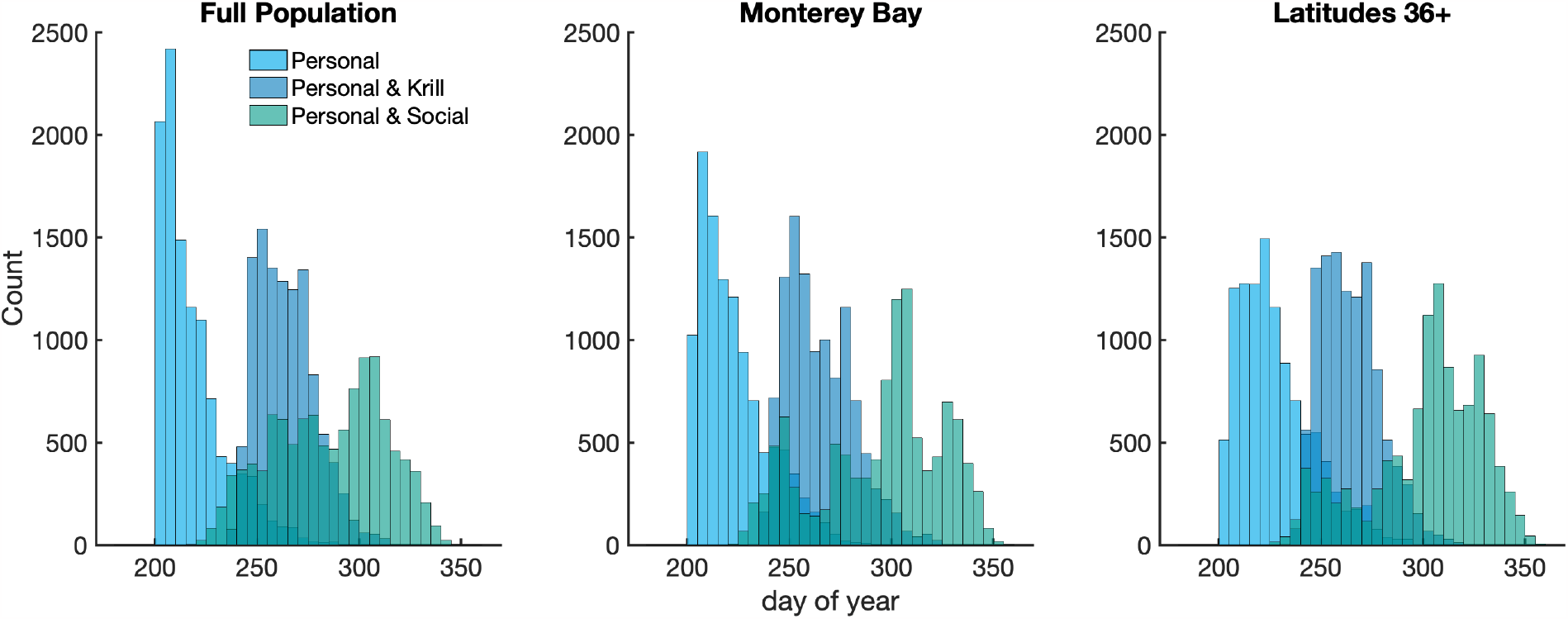
Histograms of median southward migration dates from random parameter trials. Data given for (a) the full population, (b) the portion of the population that migrated from latitudes 36-37 (Monterey Bay), and (c) the portion of the population that migrated from latitudes 36+.

## 8 Supplementary Results & Discussion

Here, we include results from the additional null-yearday and minimum krill intake migration strategies. Agent’s migration dates were recorded as the year day that they switched into S_3_. Median migration dates of all strategies are compared to the hydrophone dataset in Figure 4. As with the similar figure in the main text, median migration dates are computed for the subset of the modeled agents whose migration initiated north of Monterey Bay. A Mann-Whitney U-test (p-values in Table 5) indicates that both the set of median migration dates for the personal and personal & minimum krill strategies are statistically different from the set of hydrophone median migration dates. The migrations of the socially informed populations and null day of year are not statistically significantly different from the hydrophone data. However, the spread of the null yearday population is much narrower than the empirical hyrdophone data.

**Table 5:**
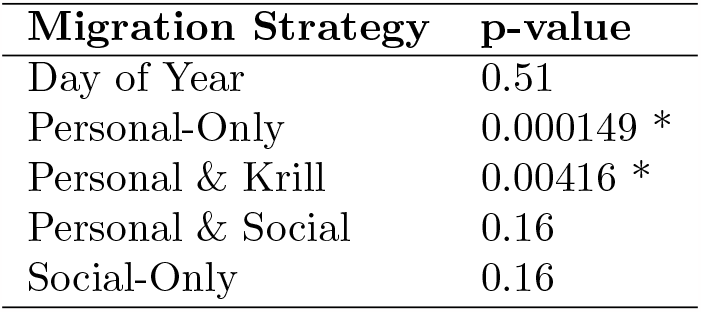
Results of the Mann-Whitney U-test to determine if distributions of median migration dates are identical to the hydrophone data. The p-values are rounded to 3 significant digits. A * indicates a statistically significant difference with a p-value *<* 0.005.

Representative yearly migration distributions are shown for the (null) yearday, personal, and personal & social strategies in Figures 5-6. The distributions for the personal & social population consistently show late-season migrations with one or more southward migrations waves.

**Figure 4.**
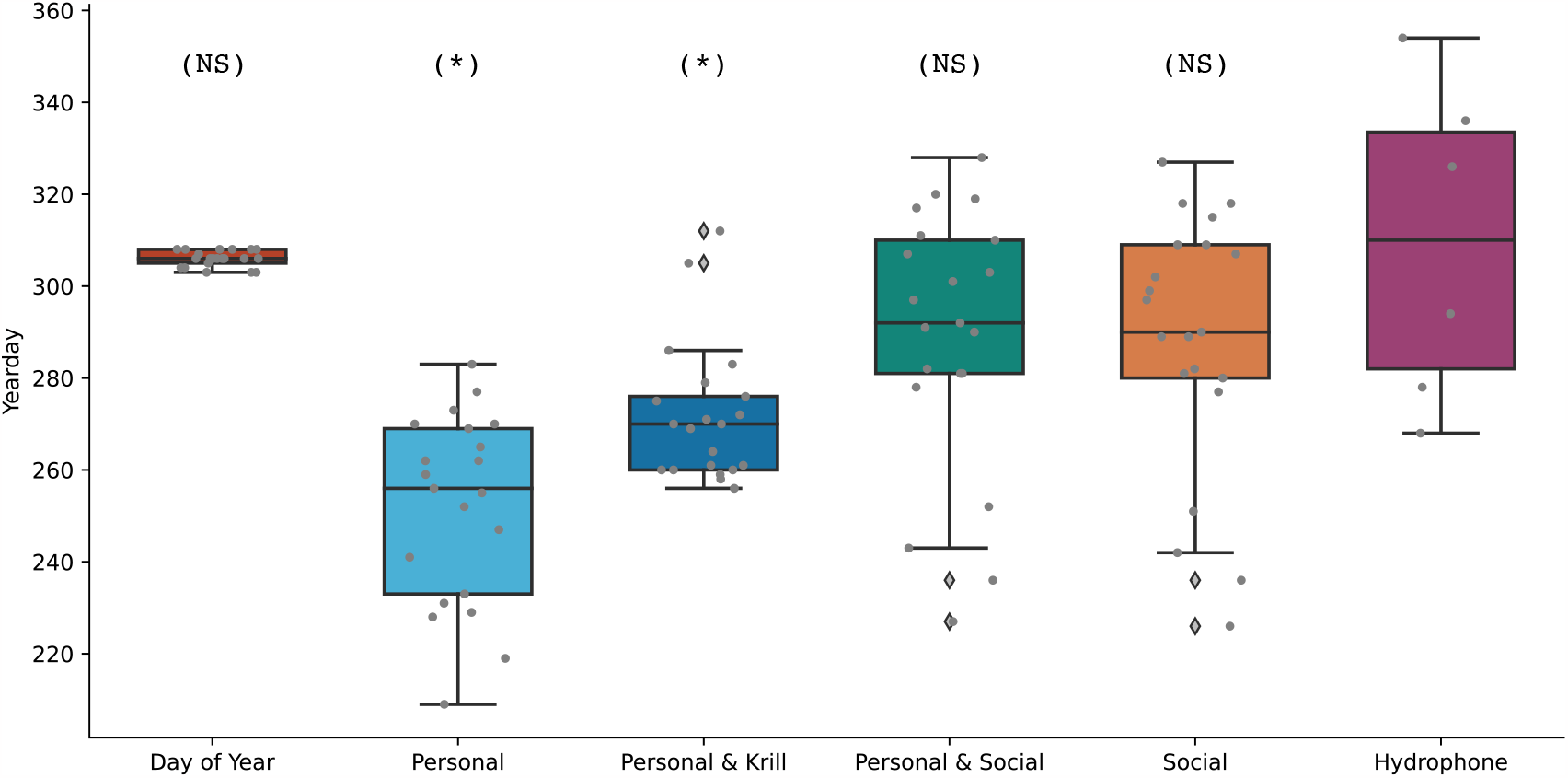
Median migration dates from MARS hydrophone data and modeled migration mechanisms. Data aggregated across all 21 years. Migration statistics for each modeled migration mechanism were calculated using the subset of the agents whose migration initiated north of Monterey Bay. Boxplots show the distribution of year median migration dates (shown in gray dots). The (*) label indicates a statistically significant difference between the set of median migration dates of the modeled mechanism and the hydrophone dataset and the (NS) label indicates no significant difference.

**Figure 5.**
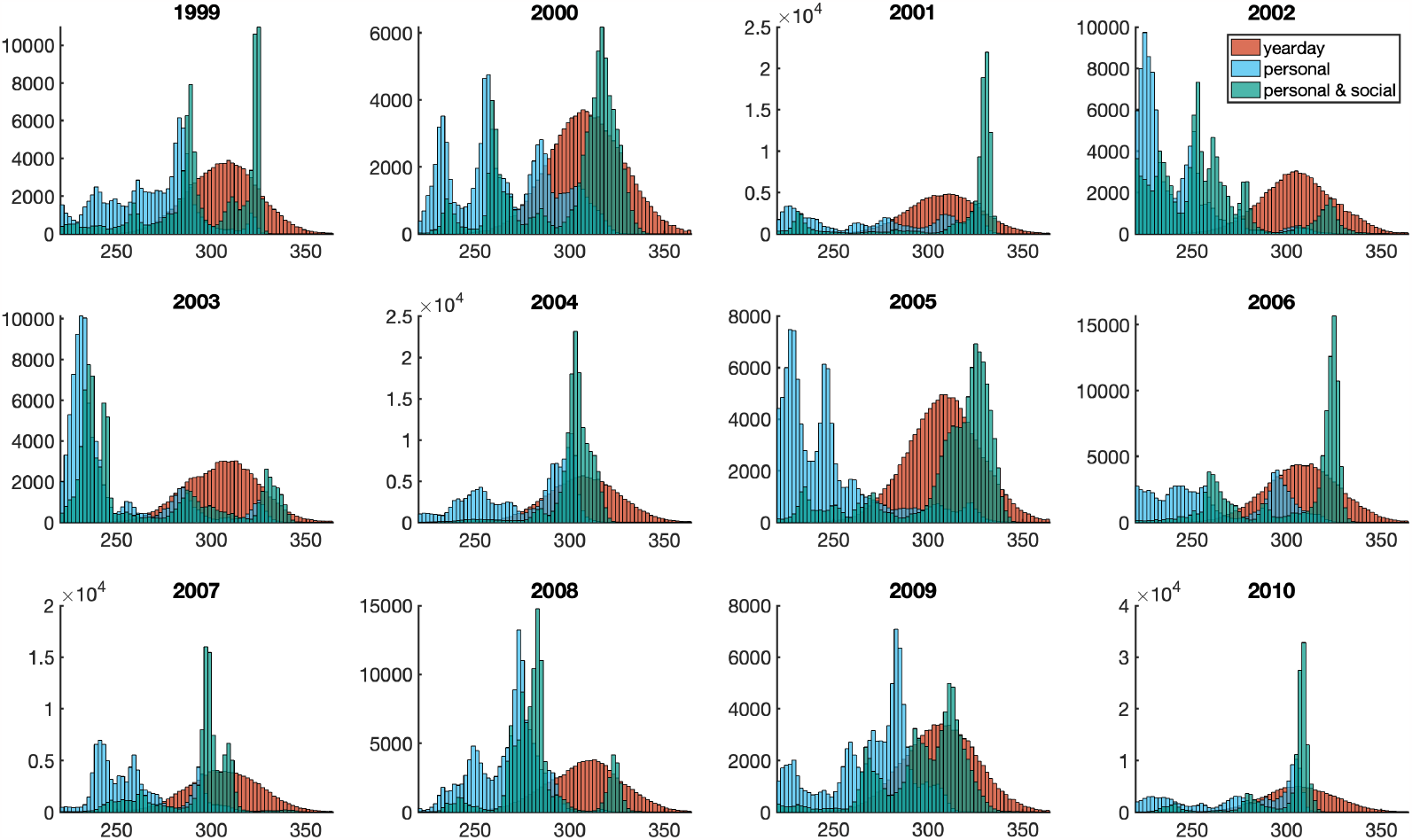
Yearly migration distributions. Representative migration distributions of personal, personal & social, and day of year migration strategies. Histograms show the yearday of migration initiation. Subset of population whose migration initiated north of Monterey Bay. Data compiled from 100 simulations for each year, with 2,000 agents per simulation.

**Figure 6.**
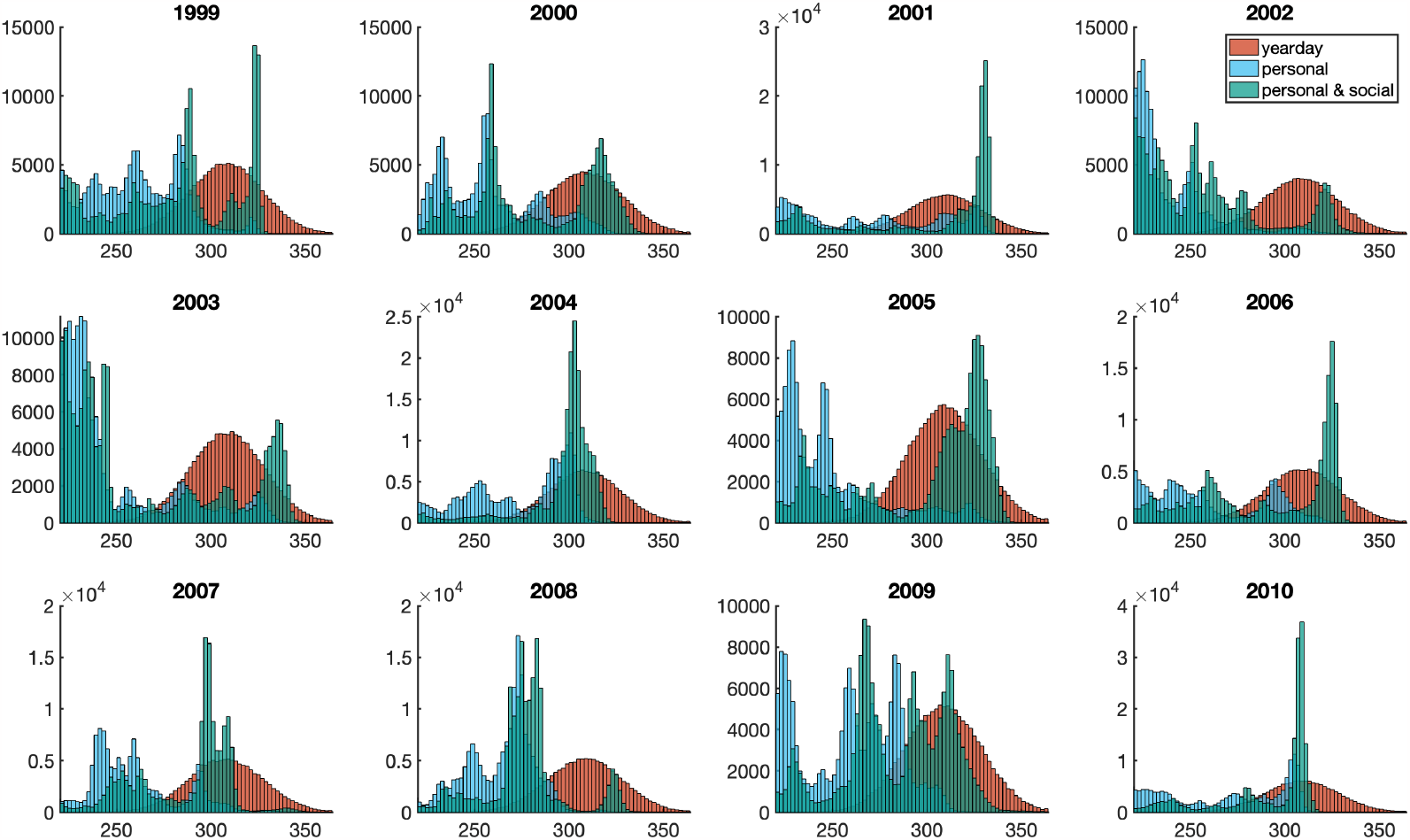
Yearly migration distributions. Representative migration distributions of personal, personal & social, and day of year migration strategies. Histograms show the yearday of migration initiation. Migration distributions shown for the full population. Data compiled from 100 simulations for each year, with 2,000 agents per simulation.

The migration dates of the null yearday population are largely consistent across yearly prey conditions and the population is unable to respond to the dynamic environment (Figure 7). The yearday population does maintain a high krill intake across the various prey conditions, largely due to the population remaining in the domain for almost the entire foraging season. The minimum krill intake requirement delays the southward migration dates (as compared to the personal population, Figures 4-7). However, the minimum prey intake also reduces the flexibility and ability to adapt to dynamic prey conditions. In years with lower than average prey, the personal & krill population has later southward migration dates as the agents continue foraging in poor conditions in an attempt to achieve the minimum prey intake.

**Figure 7.**
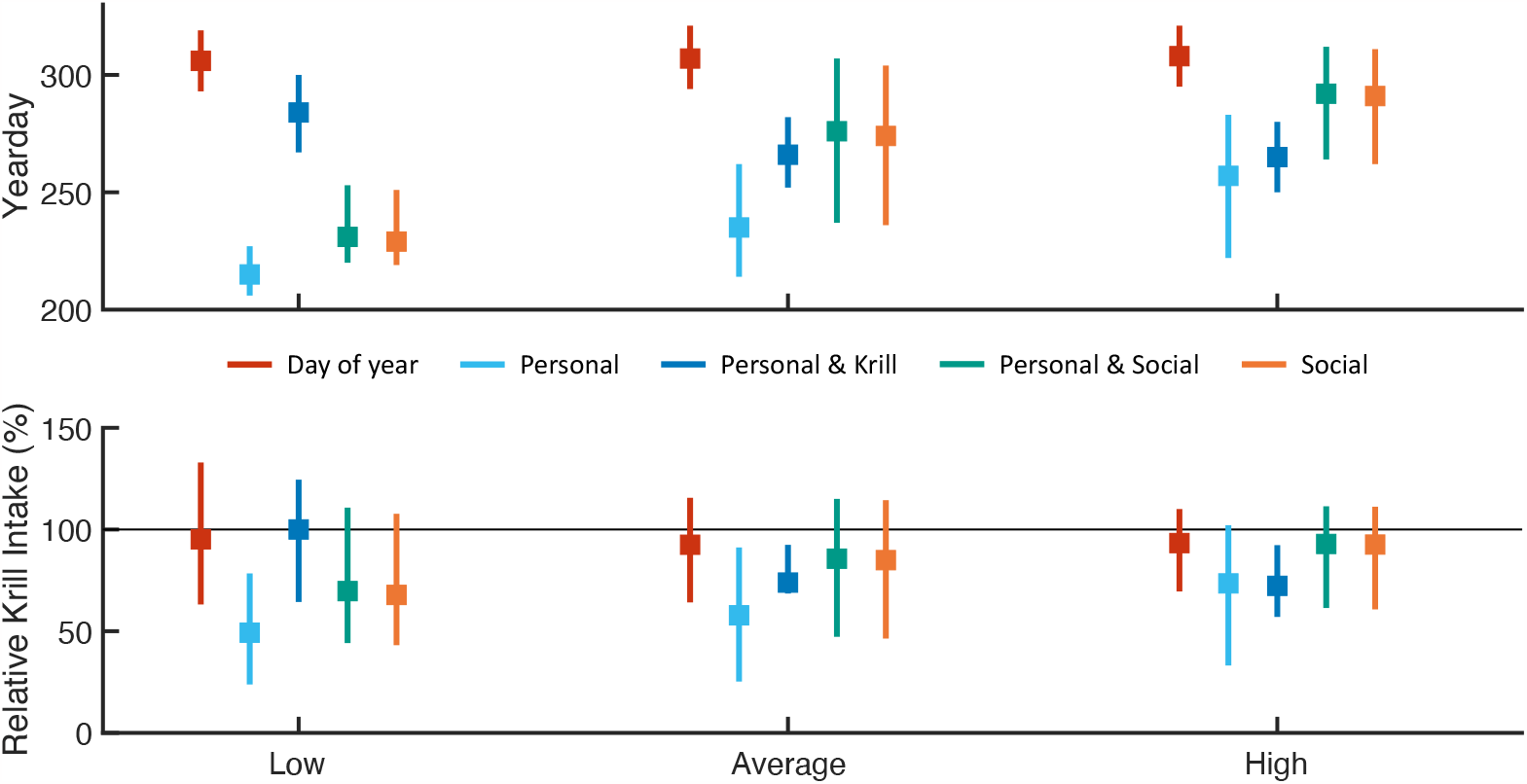
Southward migrations by krill availability. Southward migration distributions and krill intake for modeled populations separated by krill availability. Boxplots show (a) IQR of migration distributions and (b) relative krill intake for each migration mechanism. Values in (b) are computed as a percentage of the total non-migratory (null) population intake. Grey line indicates the median intake of the null population. Results from years 1990-2010 aggregated by yearly krill availability.

Figure 8 provides additional insight on the impact of the maximum call radius. Migration distributions for three years of the personal & social strategy are show in Figure 8a. Across the three years, we see convergence of the median migration dates and a narrowing of the interquartile range, indicating an increased call radius leads to a more collective migration (additionally supported by Figure 8b).

**Figure 8.**
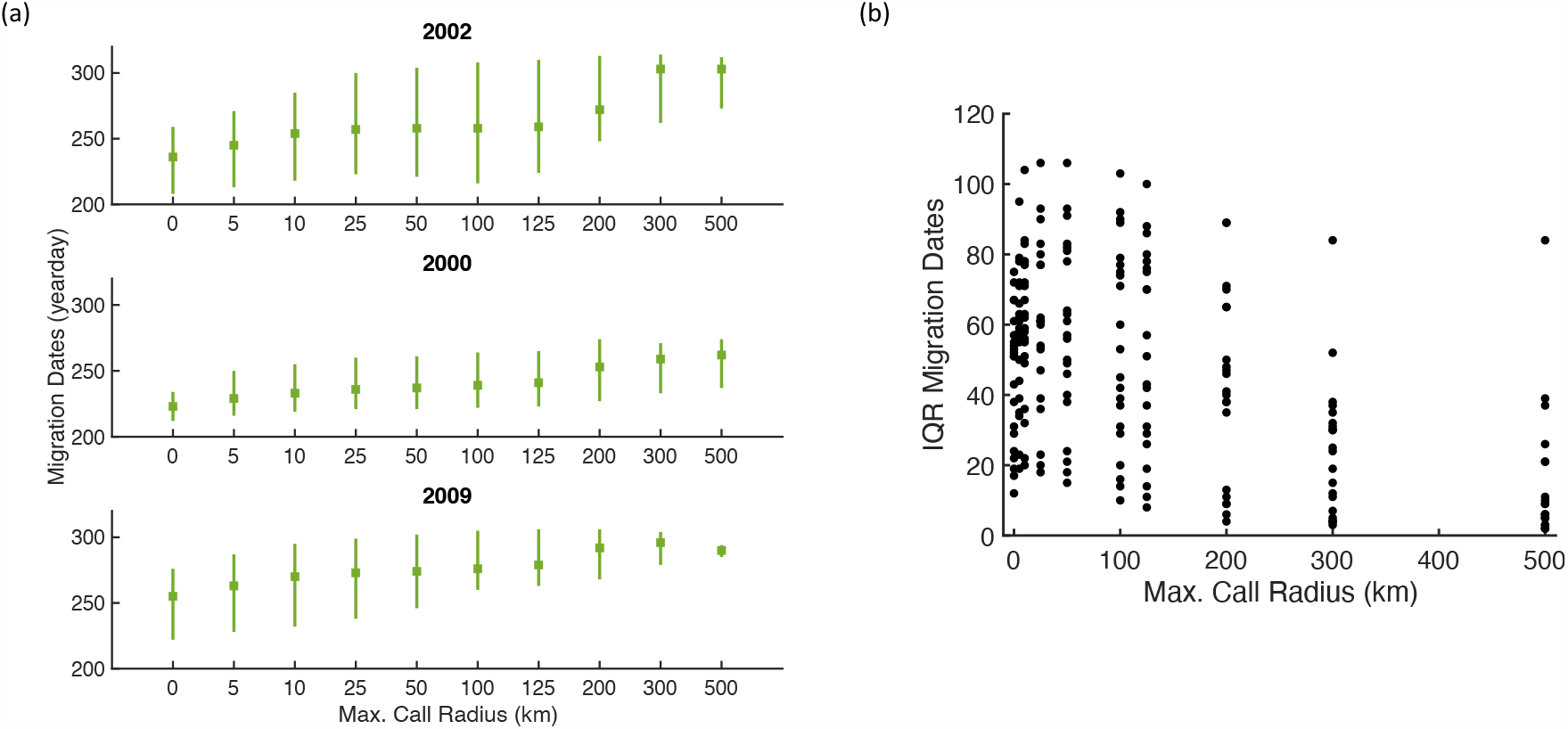
Impact of maximum call radius. (a) Representative yearly boxplots of southward migration distributions as a function of maximum call radius. (b) Interquartile range (IQR) of migration dates as a function of maximum call radius. Data from years 1990-2010. Both sets of results are from the personal & social model.

