## Supplemental Information for "Long-distance communication can enable collective migration in a dynamic seascape"

Table 1: Description of state-variables defined to each agent. Variables are unitless, unless otherwise specified. An asterisk next to the state variable name indicates a variable that is only used in models with social calls.

| State Variable | Description |
|---|---|
| Behavioral state | Value of 1-4 defining transiting and foraging behavior |
| Location | $(x, y)$-coordinate pairs, giving the distance (in meters) from the southwest corner of the domain. Continuous variable. |
| krill | Value of ROMS krill at location of agent on each time step |
| SST | Temperature value (°C; ROMS) at location on each time step |
| *Sex | Assigned male or female at random, time independent |
| *Calling behavior | Value of 0 or 1 indicates if agent is calling at current time step |
| *Received call signals | Average value of calls heard at each time step |

(4 time steps). Details of each process are included in Section 2. Movement updates are selected from state-dependent step length and turning angle distributions.

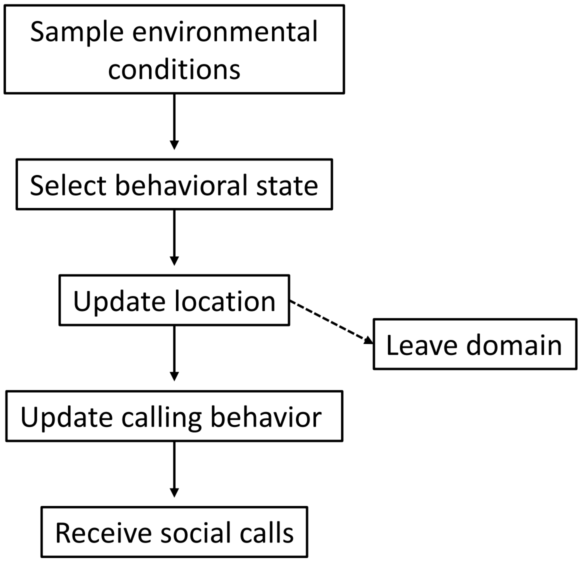


Figure 1: Schematic of underlying IBM algorithm. Process represents a single time step and is completed for each agent.

#### 2 Submodels and Model Details

This section represents the *Submodels* portion of the ODD protocol. Subsections define the processes in Figure 1.

Throughout the model description, the subscript $n$ will indicate a quantity or variable for an individual agent, where $n \in \{1, 2, \ldots, 2000\}$. Additionally, $t$ is used to represent the time step of the simulation. Model time steps are 6-hours in length and simulations are initiated on July 1 ($t_0$). The yearday $\tilde{t}$ can be extracted from the time step $t$ using the floor-function as $\tilde{t} = t_0 + \lfloor \frac{t}{4} \rfloor$. Variables are summarized in Table 2 and notation is consistent with that of the main text.

#### 2.1 Social Communication

The social communication process is divided into the call producers and call receivers.

##### 2.1.1 Call Producers

The call signal sent by agent $n$ on at time step $t$ is defined as

$$\delta_n(t) = \tanh\left[b_1\left(\bar{x}_n(t) - b_2\right)\right]$$

where $\bar{x}_n(t)$ denotes the average 24-hour foraging behavior

$$\bar{x}_n(t) = \frac{1}{4}\sum_{j=1}^{4} \mathbb{1}_{2,4}\left(s_n(t-j)\right), \qquad \mathbb{1}_{2,4}\left(s_n(t)\right) = \begin{cases} 1 & s_n(t) \in \{2,4\} \\ 0 & s_n(t) \in \{1,3\} \end{cases}.$$

Here, $\bar{x}_n(t)$ takes on a value between 0 and 1, with 0 representing all transit and 1 all forage behaviors. The call signal $\delta_n(t)$ is intended to be a time-dependent measure of foraging success analogous to the ratio $\mathrm{CI}_{\mathrm{night}}$:$\mathrm{CI}_{\mathrm{day}}$ (Oestreich et al., 2022; Oestreich et al., 2020).

At the start of the simulation, 50% of agents are randomly assigned as male (call producers) and on each time step, a subset of male whales are randomly selected to produce calls. The proportion of male whales calling on each time step increases linearly from 5% on July 1 to 30% on December 31.

##### 2.1.2 Call Receivers

Define $M^n(t)$ to be the subset of whales whose calls are heard by whale $n$ at $t$, $N_n(t) = |M_n(t)|$ (cardinality of the set $M_n(t)$) to be the number of calls received, and $R_{j,n}(t)$ the distance (in meters) between two agents $n$ and $j$. Then, the call signal received by whale $n$ at time $t$ is an average of the all calls received, weighted by the inverse-square law dependent decay of the call amplitude

$$\sigma_m(t) = \frac{1}{N_n(t)} \sum_{j \in M_n(t)} ISL_j(t)\delta_j(t), \qquad ISL_j(t) = \begin{cases} \left[A_0 - \left|20\log_{10}\left(\frac{1}{R_{j,n}(t)}\right)\right|\right] & R_{j,n}(t) < R_{\max} \\ 0 & R_{j,n}(t) \geq R_{\max} \end{cases}.$$

Call signals are capped at a maximum radius $R_{\max}$. Calling behaviors and the inverse-square law amplitude decay are independent of call radii.

| **Variable** | **Description** |
|---|---|
| $s_n(t)$ | Behavioral state of whale $n$ at time step $t$ |
| $\rho_n(t)$ | Krill density at location of whale $n$ at time step $t$ |
| $\bar{x}_n(t)$ | Daily foraging rate of whale $n$ at time step $t$ |
| $\delta_n(t)$ | Call signal sent by whale $n$ at time step $t$ |
| $\sigma_n(t)$ | Call signal received by whale $n$ at time step $t$ |
| $\bar{\omega}_n(t)$ | Foraging effectiveness of whale $n$ at time step $t$ |
| $\tau$ | State transition probability matrix |

Table 2: **Model variables.** Summary and descriptions of model variables.

| **Parameter** | **Description** | **Default Value** | **Range of Values** |
|---|---|---|---|
| **Calling Behavior** | | | |
| $b_1$ | Steepness of call signal | 8 | [3, 10] |
| $b_2$ | Call signal threshold | 0.38 | [0.3, 0.4] |
| $A_0$ | Strength of call at source | 180 dB | |
| $R_{\max}$ | Maximum call radius | 125 km | |
| **Transition to Southward Migration** | | | |
| $c_1$ | Steepness of foraging transition probability | 15 | [10, 20] |
| $c_2$ | Foraging transition threshold | 0.2 | [0.05, 0.3] |
| $c_3$ | Steepness of social call transition probability | 0.22 | [0.1, 0.25] |
| $c_4$ | Social call transition threshold | -10 | [-20, 0] |
| $\kappa_{\min}$ | Minimum krill intake threshold | 100 | |

Table 3: **Calling and migration model parameters.** Summary and descriptions of model parameters related to calling behavior and southward migration. Model results computed using default parameter values. Sensitivity analysis conducted over the listed range of values. Parameter values with no listed range were not included as part of the sensitivity analysis. Parameters without units are unitless.

#### 2.2 Selecting behavioral states

The four behavioral states $\mathcal{S}_{1,2,3,4}$ represent transiting and foraging behaviors during the northward foraging and southward breeding migrations. Each behavioral state is associated with characteristic movements defined by step length and turning angle distributions.

Transitions between the four behavioral states $\mathcal{S}_{1,2,3,4}$ are governed by the state transition probability matrix

$$\tau = \begin{pmatrix} \mathcal{S}_1 \to \mathcal{S}_1 & \mathcal{S}_1 \to \mathcal{S}_2 & \mathcal{S}_1 \to \mathcal{S}_3 & \mathcal{S}_1 \to \mathcal{S}_4 \\ \mathcal{S}_2 \to \mathcal{S}_1 & \mathcal{S}_2 \to \mathcal{S}_2 & \mathcal{S}_2 \to \mathcal{S}_3 & \mathcal{S}_2 \to \mathcal{S}_4 \\ \mathcal{S}_3 \to \mathcal{S}_1 & \mathcal{S}_3 \to \mathcal{S}_2 & \mathcal{S}_3 \to \mathcal{S}_3 & \mathcal{S}_3 \to \mathcal{S}_4 \\ \mathcal{S}_4 \to \mathcal{S}_1 & \mathcal{S}_4 \to \mathcal{S}_2 & \mathcal{S}_4 \to \mathcal{S}_3 & \mathcal{S}_4 \to \mathcal{S}_4 \end{pmatrix}$$

$$= \begin{pmatrix} p_1 & p_2 & p^* & 0 \\ p_1 & p_2 & p^* & 0 \\ 0 & 0 & p_3 & p_4 \\ 0 & 0 & p_3 & p_4 \end{pmatrix}, \qquad \begin{aligned} p_1 &= \mathbb{P}\left(s_n(t+1) = 1 | s_n(t) \in \{1,2\}\right) \\ p_2 &= \mathbb{P}\left(s_n(t+1) = 2 | s_n(t) \in \{1,2\}\right) \\ p_3 &= \mathbb{P}\left(s_n(t+1) = 3 | s_n(t) \in \{3,4\}\right) \\ p_4 &= \mathbb{P}\left(s_n(t+1) = 1 | s_n(t) \in \{3,4\}\right) \end{aligned}$$

where $s_n(t) \in \{1,2,3,4\}$ is the behavioral state of whale $n$ at time step $t$. Allowed transitions are show in the schematic in Figure 1 in the main text.

Southward migration strategies are incorporated in the southward transition probability $p^*$. Tested migration strategies are based on a combination of an agent's personal foraging behavior and received social information. Additionally, let $\bar{\omega}_n(t)$ be the average foraging behavior and $\bar{\sigma}_n(t)$ be the average received social information of agent $n$ over a period of $T = 40$ time steps (10 days). These are defined by

$$\bar{\omega}_n(t) = \frac{1}{T} \sum_{j=1}^{T} \mathbb{1}_{2,4}\left(s_n(t-j)\right) \cdot \rho_n(t-j), \qquad \bar{\sigma}_n(t) = \frac{1}{T} \sum_{j=1}^{T} \sigma_n(t-j).$$

Migration strategies are encoded in the transition probability $p^* = \mathbb{P}\left(s_n(t+1) = 3 | s_n(t) \in \{1,2\}\right)$. The transition probability functions associated with each strategy are given below. Although the subscript $n$ is omitted, all transition probabilities are assigned for each agent. All parameters are included in Table 3.

1. Individual foraging efficiency (personal):

$$p^*_{\text{per}} = \mathbb{P}\left(s_n(t+1) = 3 | s_n(t) \in \{1,2\};\ \bar{\omega}_n(t)\right) = \left[1 + \exp\left(c_1\left(\bar{\omega}_n(t) - c_2\right)\right)\right]^{-1}$$

2. Individual foraging efficiency and minimum krill intake (personal & min krill):

$$p^*_{\text{per \& krill}} = p^*_{\text{per}} \cdot p^*_{\text{krill}}$$

3. Social communication (social):

$$p^*_{\text{soc}} = \mathbb{P}\left(s_n(t+1) = 3 | s_n(t) \in \{1,2\};\ \bar{\sigma}_n(t)\right) = \left[1 + \exp\left(c_3\left(\bar{\sigma}_n(t) - c_4\right)\right)\right]^{-1}$$

4. Individual foraging efficiency and social communication (personal & social):

$$p^*_{\text{per \& soc}} = p^*_{\text{per}} \cdot p^*_{\text{soc}}$$

The personal, social, and personal & social strategies and their results are described in detail in the main text. In the Supplementary Results (Section 8), we additionally include results of a migration strategy with a minimum krill intake requirement. Specifically, agents are only able to migrate if they exceed a minimum krill threshold $\kappa_{\text{min}}$. Thus, define the function

$$p^*_{\text{krill}} = \begin{cases} 1 & \kappa_n(t) > \kappa_{\text{min}} \\ 0 & \kappa_n(t) \leq \kappa_{\text{min}} \end{cases}$$

where the cumulative krill intake $\kappa_n(t)$ is found by summing the krill density $\rho$ at the agent's foraging locations

$$\kappa_n(t) = \sum_{j=1}^{t} \mathbb{1}_{2,4}\left(s_n(t-j)\right) \cdot \rho_n(t-j).$$

State transition probabilities are computed as follows. First, $p^*$ is computed and fixed. Then, for agents in $\mathcal{S}_{1,2}$ the probabilities $p_1$ and $p_2$ are defined using the fact that the rows of the STPM sum to 1. Thus, the probabilities are set to

$$p_2 = (1 - p_*)\left[\mathbb{P}_{E,K}\left(s_n(t+1) = 2 | s_n(t) \in \{1,2\}; SST, \rho\right)\right], \qquad p_1 = 1 - p^* - p_2$$

where $\mathbb{P}_{E,K}\left(s_n(t+1) = 2 | s_n(t) \in \{1,2\}; SST, \rho\right)$ is the probability of foraging due to SST and krill density defined in (Dodson et al., 2020). Parameter values for the forage-transit selection process are identical to those in (Dodson et al., 2020). Transition probability functions between $\mathcal{S}_1$ and $\mathcal{S}_2$ are identical across all presented models.

For agents in $\mathcal{S}_{3,4}$, we likewise define the probability of foraging and utilize that $p_3 + p_4 = 1$. Thus,

$$p_4 = \mathbb{P}_{E,K}\left(s_n(t+1) = 4 | s_n(t) \in \{3,4\}; SST, \rho\right)$$
$$p_3 = 1 - p_4$$

where $p_4$ has a stricter foraging threshold (high krill density required for foraging). Transition probabilities between $\mathcal{S}_3$ and $\mathcal{S}_4$ are identical across all presented models that include the southward behavioral states.

##### 2.2.1 Sampling Environmental Conditions and Location Updates

On each time step, agents sample and store as state variables the SST and krill density at their current location (values from ROMS data). These environmental parameters are used to compute the elements of the state transition probability matrix.

Regardless of migration strategy, movement updates are selected from distinct turning angle and step length distributions associated with characteristic transit and forage behaviors. See Dodson et al., 2020 for movement distributions.

### 3 Model Assumptions

Here, we summarize and clarify model assumptions.

- Behavioral states and positions are updated every six hours. Transition probabilities within the north and southward migration categories (between states $\mathcal{S}_1 \to \mathcal{S}_2$ and $\mathcal{S}_3 \to \mathcal{S}_4$) are based only on SST and prey levels.
- Transition probabilities between $\mathcal{S}_1 \to \mathcal{S}_2$ are identical across all presented models.
- Transition probabilities between $\mathcal{S}_3 \to \mathcal{S}_4$ are identical across all presented models that include the southward behavioral states.
- Individuals commit to southward migration and are not permitted to transition from states $\mathcal{S}_{3,4}$ to $\mathcal{S}_{1,2}$.
- Agents are not bound to the domain and will freely leave if their movement updates take them outside the domain.

| Krill Availability | Years |
|---|---|
| Low | 1990, 1992, 1997, 2002 |
| Average | 1993, 1994,1998, 1999, 2000, 2003, 2005, 2006, 2007, 2009 |
| High | 1991, 1995, 1996, 2001, 2004, 2008, 2010 |

Table 4: Classification of years based on krill intake of the null non-migratory population.

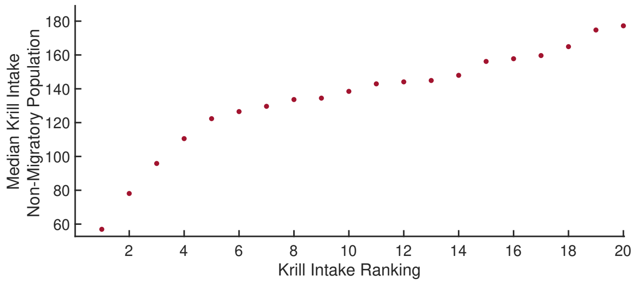


Figure 2: **Yearly krill intake.** Ranked median yearly krill intake of the null non-migratory population.

### 7 Sensitivity analysis

Robustness of the timing of the migration mechanism was tested with using random parameter samples. Realistic, but large intervals were set for all parameters (Table 3). A total of 1,000 trials were run for each year in 2000-2010 with parameters randomly selected from the set parameter ranges using Latin Hypercube Sampling procedures (Marino et al., 2008). Figure 3 shows the median migration dates for all random sample trials. Tested parameter intervals were consistent across all strategies.

The addition of social calls leads to late season migrations over the full range of realistic parameters. Notably, even the lowest median migrate dates for social strategies are within reason. The asocial strategies consistently lead to early migrations over the broad range of parameters with median dates as early as September - even when restricted to the subset of the population that migrates from the higher latitudes. The results of this sensitivity analysis are important - they show that including social information in migration decisions leads to a late season migration over a broad range of parameters and conditions.

The default set of parameters stated in Table 3 and used for the results in the main text were selected to give later season migrations in the personal-only population (migration timings in the upper tails of the distributions in Figure 3).

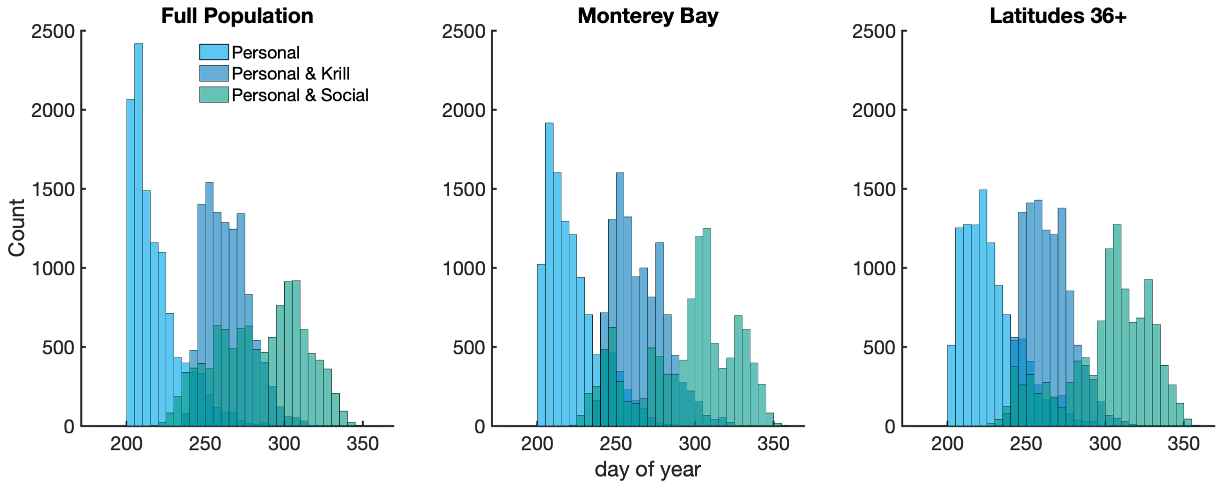


Figure 3: **Histograms of median southward migration dates from random parameter trials.** Data given for (a) the full population, (b) the portion of the population that migrated from latitudes 36-37 (Monterey Bay), and (c) the portion of the population that migrated from latitudes 36+.

### 8 Supplementary Results & Discussion

Here, we include results from the additional null-yearday and minimum krill intake migration strategies. Agent's migration dates were recorded as the year day that they switched into $\mathcal{S}_3$. Median migration dates of all strategies are compared to the hydrophone dataset in Figure 4. As with the similar figure in the main text, median migration dates are computed for the subset of the modeled agents whose migration initiated north of Monterey Bay. A Mann-Whitney U-test (p-values in Table 5) indicates that both the set of median migration dates for the personal and personal & minimum krill strategies are statistically different from the set of hydrophone median migration dates. The migrations of the socially informed populations and null day of year are not statistically significantly different from the hydrophone data. However, the spread of the null yearday population is much narrower than the empirical hyrdophone data.

| **Migration Strategy** | **p-value** |
|---|---|
| Day of Year | 0.51 |
| Personal-Only | 0.000149 * |
| Personal & Krill | 0.00416 * |
| Personal & Social | 0.16 |
| Social-Only | 0.16 |

Table 5: Results of the Mann-Whitney U-test to determine if distributions of median migration dates are identical to the hydrophone data. The p-values are rounded to 3 significant digits. A * indicates a statistically significant difference with a p-value $< 0.005$.

Representative yearly migration distributions are shown for the (null) yearday, personal, and personal & social strategies in Figures 5-6. The distributions for the personal & social population consistently show

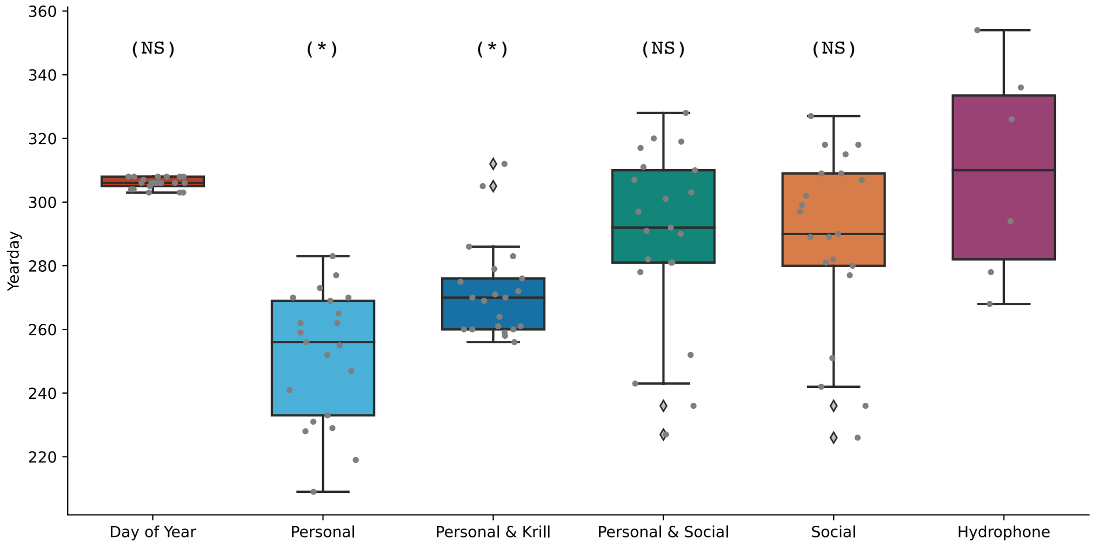


Figure 4: **Median migration dates from MARS hydrophone data and modeled migration mechanisms.** Data aggregated across all 21 years. Migration statistics for each modeled migration mechanism were calculated using the subset of the agents whose migration initiated north of Monterey Bay. Boxplots show the distribution of year median migration dates (shown in gray dots). The (*) label indicates a statistically significant difference between the set of median migration dates of the modeled mechanism and the hydrophone dataset and the (NS) label indicates no significant difference.

late-season migrations with one or more southward migrations waves.

The migration dates of the null yearday population are largely consistent across yearly prey conditions and the population is unable to respond to the dynamic environment (Figure 7). The yearday population does maintain a high krill intake across the various prey conditions, largely due to the population remaining in the domain for almost the entire foraging season. The minimum krill intake requirement delays the southward migration dates (as compared to the personal population, Figures 4-7). However, the minimum prey intake also reduces the flexibility and ability to adapt to dynamic prey conditions. In years with lower than average prey, the personal & krill population has later southward migration dates as the agents continue foraging in poor conditions in an attempt to achieve the minimum prey intake.

Figure 8 provides additional insight on the impact of the maximum call radius. Migration distributions for three years of the personal & social strategy are show in Figure 8a. Across the three years, we see convergence of the median migration dates and a narrowing of the interquartile range, indicating an increased call radius leads to a more collective migration (additionally supported by Figure 8b).

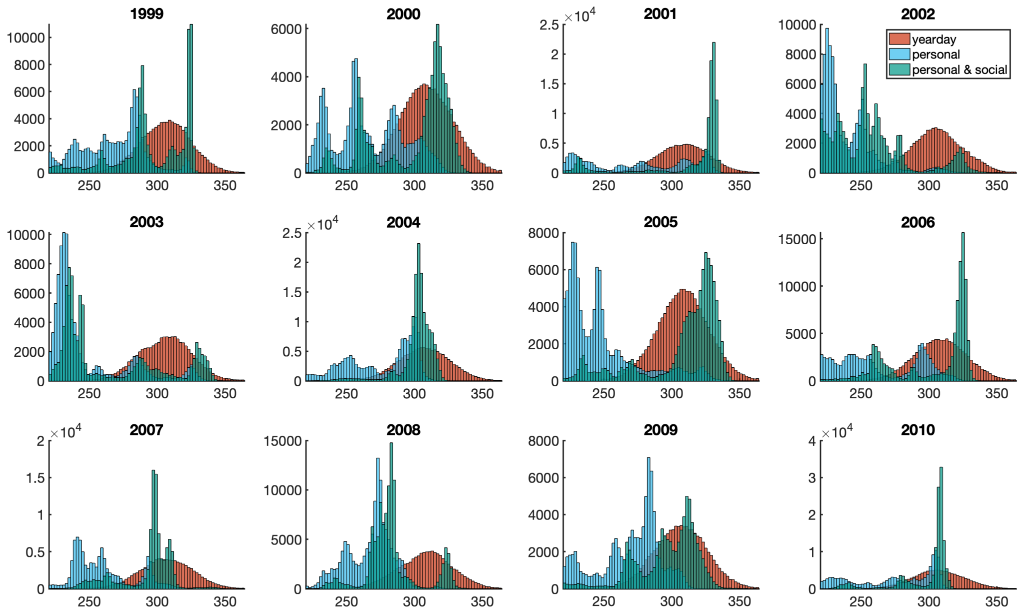


Figure 5: **Yearly migration distributions.** Representative migration distributions of personal, personal & social, and day of year migration strategies. Histograms show the yearday of migration initiation. Subset of population whose migration initiated north of Monterey Bay. Data compiled from 100 simulations for each year, with 2,000 agents per simulation.

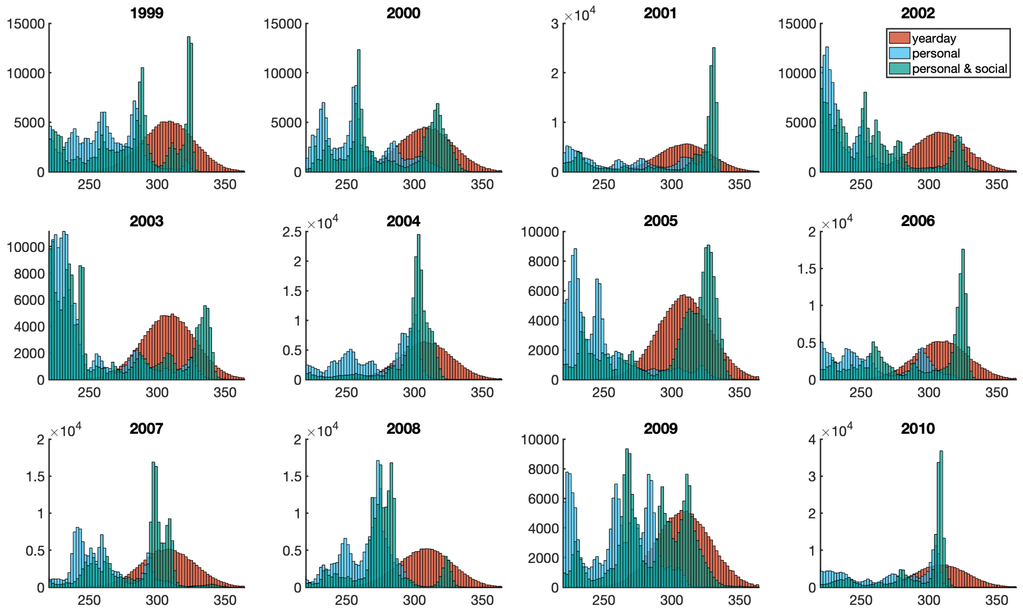


Figure 6: **Yearly migration distributions.** Representative migration distributions of personal, personal & social, and day of year migration strategies. Histograms show the yearday of migration initiation. Migration distributions shown for the full population. Data compiled from 100 simulations for each year, with 2,000 agents per simulation.

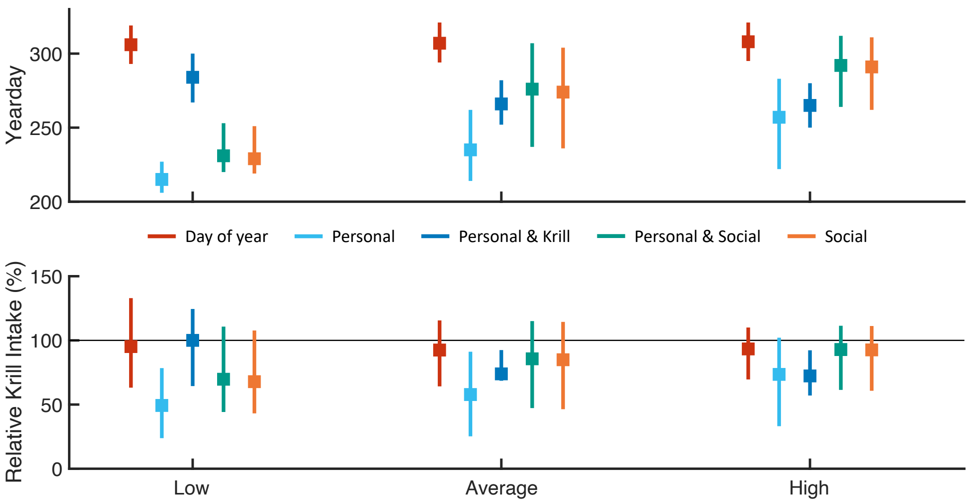


Figure 7: **Southward migrations by krill availability.** Southward migration distributions and krill intake for modeled populations separated by krill availability. Boxplots show (a) IQR of migration distributions and (b) relative krill intake for each migration mechanism. Values in (b) are computed as a percentage of the total non-migratory (null) population intake. Grey line indicates the median intake of the null population. Results from years 1990-2010 aggregated by yearly krill availability.

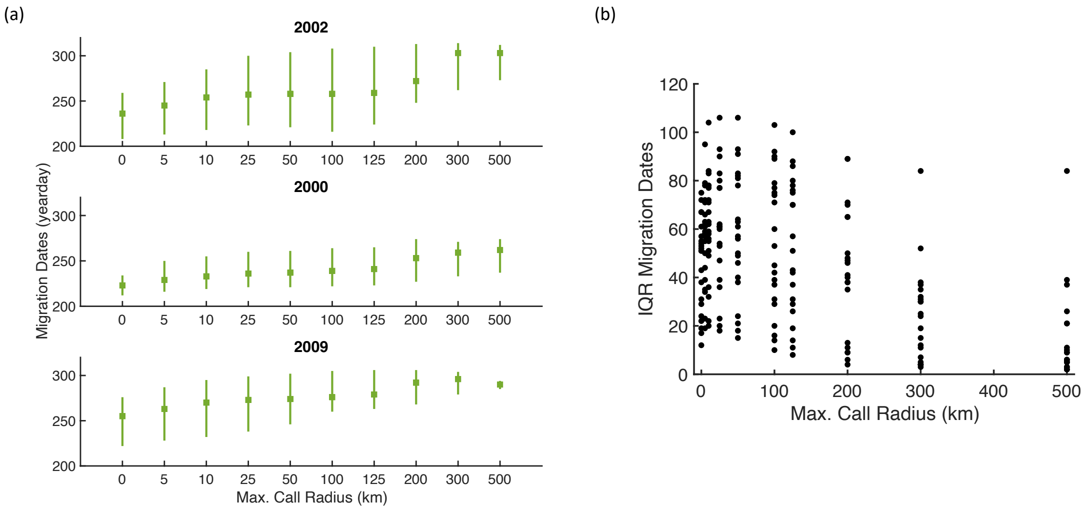


Figure 8: **Impact of maximum call radius.** (a) Representative yearly boxplots of southward migration distributions as a function of maximum call radius. (b) Interquartile range (IQR) of migration dates as a function of maximum call radius. Data from years 1990-2010. Both sets of results are from the personal & social model.
